# Single-nucleus transcriptomics reveals cell type-specific and time-dependent effects of psilocybin and ketamine on gene expression

**DOI:** 10.1101/2025.01.04.631335

**Authors:** Clara Liao, Ethan O’Farrell, Amanda M. Weiner, Yaman Qalieh, Neil K. Savalia, Matthew J. Girgenti, Kenneth Y. Kwan, Alex C. Kwan

## Abstract

There is growing interest to investigate classic psychedelics and ketamine as therapeutics for mental illnesses. Previous studies have demonstrated that one dose of psilocybin or ketamine leads to persisting neural and behavioral changes. The durability of these effects suggests that there are likely alterations in gene expression at the transcriptional level. In this study, we performed single-nucleus RNA sequencing of the dorsal medial frontal cortex of male and female mice. Samples were collected at 1, 2, 4, 24, or 72 hours after psilocybin or ketamine administration and from control animals. At baseline, major subtypes of excitatory and GABAergic neurons selectively express particular serotonin receptor transcripts. The psilocybin-evoked differentially expressed genes in excitatory neurons are involved in synaptic plasticity, distinct from genes enriched in GABAergic neurons, which contribute to mitochondrial function and cellular metabolism, and non-neuronal glial cells. The effect of psilocybin on gene expression is time-dependent, including an early phase at 1 hour followed by a late phase at 72 hours of transcriptional response after administration, and differs from the changes following ketamine administration, which peaks at 2 – 4 hours. Collectively, the results provide a resource for understanding the cell type-specific and time-dependent changes in gene expression induced by psilocybin and ketamine in the mouse medial frontal cortex, which may underpin the drug’s long-term effects on neural circuits and behavior.

## INTRODUCTION

Psilocybin is a serotonergic psychedelic that has shown promise as a potential therapeutic for mental health conditions. Data from several clinical trials indicate that psilocybin with psychological support can relieve symptoms in patients with major depressive disorder or treatment-resistant depression^1, 2^. Ketamine has long been studied for its rapid antidepressant effects^3^, paving the way for FDA approval of intranasal esketamine for treatment-resistant depression in the United States^4^. Although the neurobiological basis for the long-term effects of psilocybin and ketamine is not understood, there is growing evidence that the antidepressant effect may relate to the drug-evoked structural neural plasticity in the prefrontal cortex^5^. Numerous studies have demonstrated that ketamine promotes the formation of dendritic spines in cortical pyramidal neurons^6–8^. Similarly, psilocybin elevates the density of dendritic spines in the mouse medial frontal cortex, an effect that persists for over a month^9, 10^.

Long-lasting modifications to synaptic connections and behavior require changes in gene expression^11, 12^. Indeed, ketamine has been shown to produce sustained transcriptional responses in mice^13, 14^. For psychedelics, less is known, but previous studies of mRNA transcript and protein levels identified differential expression of genes for synaptic function, glutamate receptor signaling, and immune response in the neocortex after a single dose of a classic psychedelic such as 2,5-dimethoxy-4-iodoamphetamine (DOI)^15–18^, lysergic acid diethylamide (LSD)^15, 19^, 5-methoxy-*N*,*N*-dimethyltryptamine (5-MeO-DMT)^20^, or psilocybin^18, 21, 22^. Although these results demonstrated transcriptional alterations after drug exposure, most early studies relied on methods such as quantitative PCR and microarrays, which were limited by their focus on preselected candidate genes. A genome-wide transcriptomic investigation can yield novel insights, as exemplified by recent studies of transcriptional responses to drugs such as cocaine^23–25^, heroin^26^, fentanyl^27^, DOI^17^, MDMA^28^, as well as in mouse models of chronic stress^29, 30^ and early life stress^31, 32^.

A comprehensive understanding of the transcriptional responses requires not only identifying genes but also understanding the temporal dynamics of their regulation. This is particularly important for psychedelic and dissociative compounds, where biological effects unfold across markedly distinct timescales. In humans, the oral administration of psilocybin induces subjective effects that peak by as early as 60 minutes and wane by 3 – 6 hours, coinciding with changes in plasma psilocin levels^33^. Long after the subjective effects wear off, therapeutic effects for major depression can be detected with an onset at 2 – 8 days^1, 2^, and persist for at least 6 weeks after the initial administration^2^. Ketamine exhibits similar acute and sustained effects. Dissociative and perceptual effects are prominent during the first hour after administration^3^, whereas antidepressant effects emerge within 4 hours and persist for a few weeks thereafter^3, 4^. Therefore, efforts to understand psilocybin- and ketamine-evoked changes in gene expression should consider time as a key parameter.

To create a resource for investigating the transcriptional effects of psilocybin and ketamine, we performed single-nucleus RNA sequencing (snRNA-seq) of the dorsal medial frontal cortex from male and female mice after drug administration. To capture both acute and long-term changes, we collected tissues at 1, 2, 4, 24, and 72 hours after psilocybin or ketamine exposure, as well as from control animals. We delineated changes in gene expression due to psilocybin or ketamine administration in 16 major excitatory neurons, GABAergic inhibitory neurons, and non-neuronal cell types. To facilitate exploration and further analysis of these data, we have made the complete dataset publicly available through an interactive web portal (https://psilo-seq.kwanlab.org/). Collectively, this resource provides a foundation for investigating the cell type-specific and time-dependent changes to the transcriptional landscape in the mouse medial frontal cortex after a single dose of psilocybin or ketamine.

## RESULTS

### Single-nucleus RNA sequencing of the mouse medial frontal cortex at various timepoints after a single dose of psilocybin

In this study, we focus on the mouse dorsal medial frontal cortex, which encompasses the anterior cingulate cortex (ACAd) and the premotor cortex (medial MOs). This region is important for the drug action of psilocybin and ketamine, showing robust c-Fos response in a whole-brain mapping study^34^, corroborating reports of drug-evoked structural plasticity after psilocybin^9^ or ketamine^7^. We administered C57BL/6J mice with psilocybin (1 mg/kg, i.p.) or ketamine (10 mg/kg, i.p.), and then collected tissue corresponding to the dorsal medial frontal cortex of both hemispheres via microdissection at 1, 2, 4, 24, or 72 hours after drug administration (**Fig. 1a**). Additionally, we collected control tissue from animals that received no injection. The tissue was processed through nuclear dissociation, nuclei sorting, and barcoding for snRNA-seq via the 10x Genomics Chromium Single Cell 3’ v3 platform. We planned for and processed samples from 49 animals, including 4 for each time point and 9 controls, but excluded 4 animals due to incorrect sequencing depth specification and 3 animals due to low number or low-quality nuclei (see **Methods; Supplementary Table 1**). For the remaining 42 samples, we started with 145,593 cells for the psilocybin group, 139,628 cells for the ketamine group, and 67,600 cells for the controls. We targeted the sequencing to 25,000 reads per nucleus for each sample. Initial processing confirmed an average of 29,562 reads per nucleus. We removed cells that did not meet quality control criteria (see **Methods**). The final dataset consisted of 133,564 cells for the psilocybin group (range = 20,326–33,939 single nuclei per time point, 5,818–9,631 per animal; n = 17 animals), 127,289 cells for the ketamine group (22,511–29,554 per time point, 5,991–9,478 per animal; n = 17 animals), and 61,379 cells for the control group (6,882–8,680 per animal; n = 8 animals) (**Supplementary Table 2**).

**Fig. 1:**
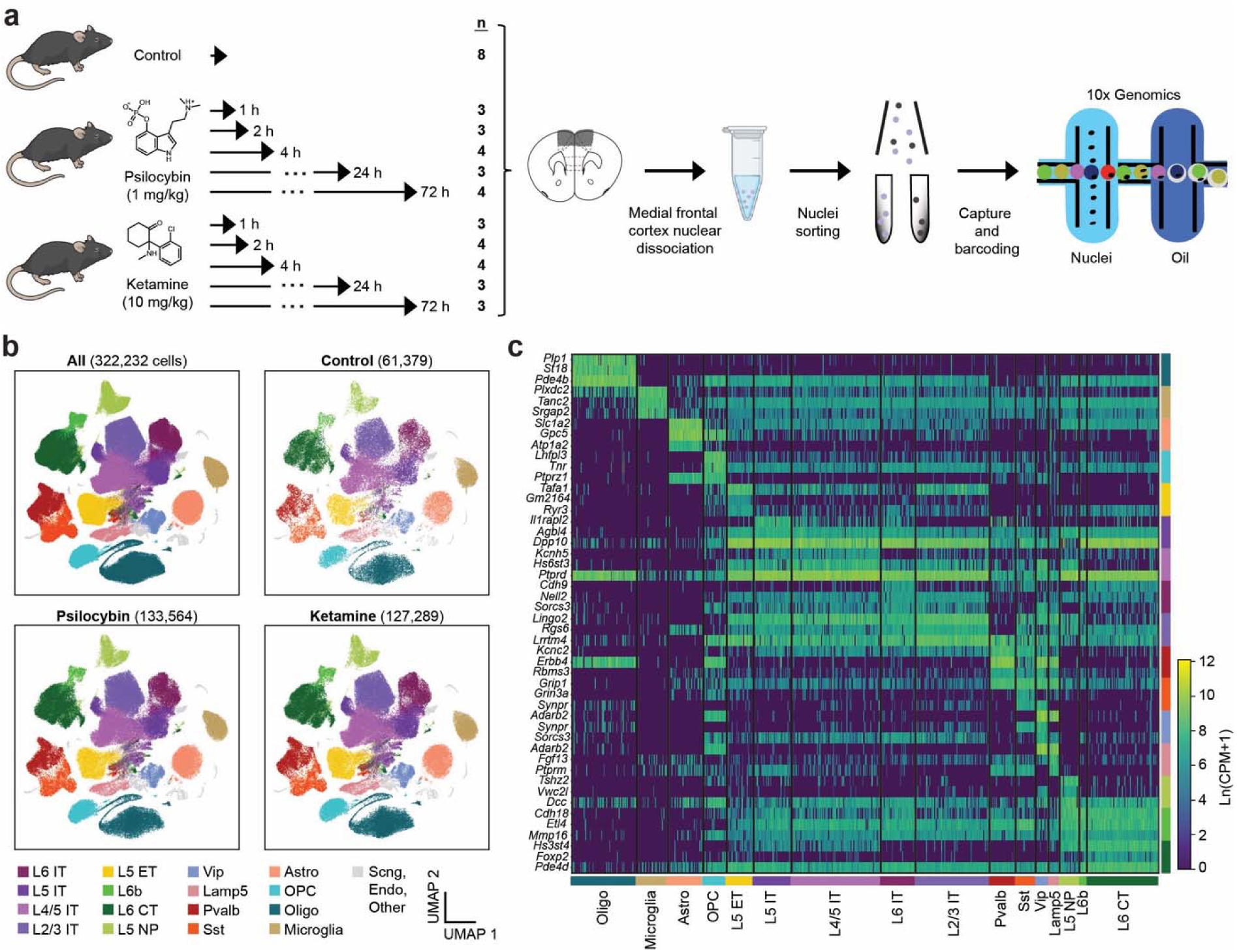
Single-nucleus RNA sequencing of mouse medial frontal cortex after psilocybin administration. **(a)** Experimental workflow. n, the number of samples for each condition. **(b)** UMAP representations of the snRNA-seq dataset, including all samples (top left), control samples only (top right), psilocybin samples only (bottom left), or ketamine samples only (bottom right). Color denotes the cell type. **(c)** Heatmap for the expression of putative marker genes for the cell types. CPM, count per million. L6 IT, layer 6 intratelencephalic neurons. L5 IT, layer 5 intratelencephalic neurons. L4/5 IT, layer 4/5 intratelencephalic neurons. L2/3 IT, layer 2/3 intratelencephalic neurons. L5 ET, layer 5 extratelencephalic neurons. L6b, layer 6b neurons. L6 CT, layer 6 corticothalamic neurons. L5 NP, layer 5 near-projecting neurons. Vip, vasointestinal peptide-expressing interneurons. Lamp5, *Lamp5*-expressing interneurons. Pvalb, parvalbumin-expressing interneurons. Sst, somatostatin-expressing interneurons. Astro, astrocytes. OPC, oligodendrocyte progenitor cells. Oligo, oligodendrocytes. Microglia, microglia. Sncg, *Sncg*-expressing interneurons. Endo, endothelial cells. Other, other annotated cell types.

We annotated cells based on the Allen Institute mouse brain taxonomy using the MapMyCells tool^35^, because the standardized cell types enabled interoperability with other datasets (**Fig. 1b**; see **Methods**). We focused on 16 major neocortical cell types. Nearly all cells were classified into one of these clusters, including 129,678 cells (97.1%) in the psilocybin group, 123,593 cells (97.1%) in the ketamine group, and 59,256 cells (96.5%) in the control group. The 16 cell types included 8 classes of excitatory neurons: layer 6 intratelencephalic neurons (004 L6 IT CTX Glut; 5.9% of classified cells across all samples), layer 5 intratelencephalic neurons (005 L5 IT CTX Glut; 6.5%), layer 4/5 intratelencephalic neurons (006 L4/5 IT CTX Glut; 15.2%), layer 2/3 intratelencephalic neurons (007 L2/3 IT CTX Glut; 12.6%), layer 5 extratelencephalic neurons (022 L5 ET CTX Glut; 4.6%), layer 6b neurons (029 L6b CTX Glut; 1.3%), layer 6 corticothalamic neurons (029 L6 CT CTX Glut; 12.3%), and layer 5 near-projecting neurons (L5 NP CTX Glut; 3.4%). In addition, there were 4 classes of GABAergic neurons: vasoactive intestinal peptide-expressing interneurons (046 Vip Gaba; 2.2%), *Lamp5*-expressing interneurons (049 Lamp5 Gaba; 1.9%), parvalbumin-expressing interneurons (052 Pvalb Gaba; 4.4%), and somatostatin-expressing interneurons (053 Sst Gaba; 3.4%), as well as 4 classes of non-neuronal cells: astrocytes (319 Astro-TE NN; 6.0%), oligodendrocyte progenitor cells (326 OPC NN; 4.0%) oligodendrocytes (327 Oligo NN; 11.1%), and microglia (334 Microglia NN, 5.3%). In total, 61.7%, 11.9%, and 26.4% of the analyzed cells were excitatory, GABAergic, and non-neuronal cells, respectively (**Supplementary Table 1**). As expected, these cell types displayed distinct expression profiles (**Fig. 1c**). The same cell-type clusters were detected in samples collected at different time points after psilocybin administration, ketamine administration, and in controls (**Supplementary Fig. 1**). Most cells were classified with near-perfect confidence (**Supplementary Fig. 2**), and similar groupings could be identified using an unsupervised clustering approach (**Supplementary Fig. 3**).

To facilitate access and exploration of the dataset, we developed an interactive web portal (https://psilo-seq.kwanlab.org/) that allows users to visualize the data and download either selected subsets or the entire dataset for independent analyses. While the resource includes both psilocybin and ketamine conditions, we will focus the subsequent analyses primarily on psilocybin to illustrate the biological insights that can be derived from this resource.

### Cell type-specific expression of serotonin receptor transcripts in the mouse medial frontal cortex

Following administration, psilocybin is rapidly metabolized to psilocin, which enters the brain. Psilocin binds not only to the 5-HT_2A_ receptor, but also to other serotonin receptor subtypes expressed in the neocortex, including the 5-HT_1A_ and 5-HT_2C_ receptors^36, 37^. We visualized the presence of the *Htr1a*, *Htr2a*, and *Htr2c* transcripts which encode the 5-HT_1A_, 5-HT_2A_, and 5-HT_2C_ receptors respectively in the control samples in our snRNA-seq dataset (**Fig. 2a**). The expression patterns of the three transcripts clearly differed, prompting us to quantify how the transcript levels vary across the 16 cell types (**Fig. 2b, c**). Because our cell types were annotated based on Allen Institute mouse brain taxonomy, the results can be easily compared with many other existing datasets. To illustrate interoperability, we extracted SMART-Seq single-cell sequencing data from the Allen Institute^38^, focusing on cells residing in the mouse frontal cortex, including the ACA, ALM, ORB, and PL-ILA regions (**Fig. 2d**). There was strong agreement between the two datasets. The IT and ET excitatory cell types express a high level of *Htr2a* as well as moderate amounts of *Htr1a* and *Htr2c*. Interestingly, there is also an abundance of *Htr2a* transcripts in the deep-lying layer 6b neurons, which are known for their excitatory effects on apical dendrites and higher-order thalamus^39^. Unlike excitatory neurons, the expression profile is more selective in the GABAergic cell subtypes. The Lamp5 and Pvalb subtypes express *Htr2a*, whereas the Vip subpopulation contains *Htr2c*. Although serotonin receptors have been reported in microglia^40, 41^ and astrocytes^42^, the current data suggest that *Htr* transcript levels are relatively low in the non-neuronal cells. Altogether, these analyses reveal a highly selective pattern of serotonin receptor expression in cortical cell types. Because psilocin is a non-selective serotonergic agonist that binds to a range of receptor subtypes, we anticipate that the drug can directly influence the transcriptional programs of most major excitatory and GABAergic cell types, while indirectly affecting other cell populations.

**Fig. 2:**
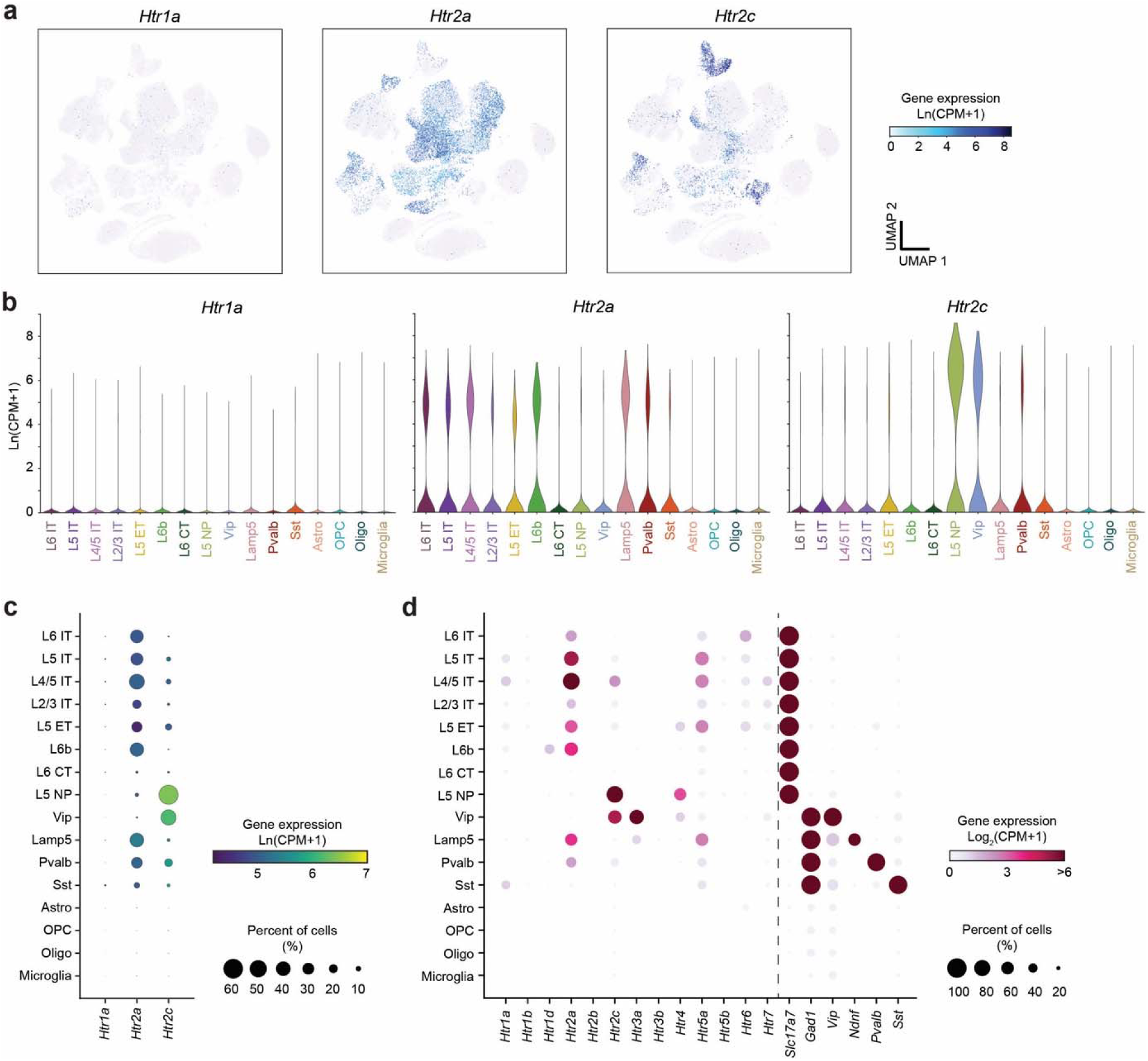
Cell type-specific expression of serotonin receptor transcripts in the mouse medial frontal cortex. **(a)** UMAP representation of the snRNA-seq dataset. Color denotes expression level (CPM, counts per million) of *Htr1a* (left), *Htr2a* (middle), or *Htr2c* (right). **(b)** The expression levels of *Htr1a* (left), *Htr2a* (middle), and *Htr2c* (right) in the 16 cell types. **(c)** Transcriptomic profiles from control mice in this study. Circle size denotes the percentage of cells with detectable expression of a gene (>0 reads). Circle color denotes expression level. **(d)** Similar to (c), but using data from the SMART-Seq dataset from Allen Institute. Single-cell transcript counts were summed from cells in the mouse ACA, ALM, ORB, and PL-ILA regions. *Slc17a7* and *Gad1* are markers of excitatory and GABAergic neurons, respectively., *Vip*, *Ndnf*, *Pvalb*, and *Sst* are markers of subtypes of GABAergic neurons.

### Psilocybin-induced transcriptional changes in frontal cortical excitatory neurons are involved in synaptic plasticity

To identify differentially expressed genes (DEGs), we used pseudobulk analysis where sequence reads were summed for each cell type and biological replicate (see **Methods**). Pseudobulk analysis has been shown to yield reliable quantifications with fewer false positives^43, 44^. We first report DEGs in the four populations of intratelencephalic excitatory neurons **(Fig. 3a, b).** Across all four IT cell types, we detected the highest number of DEGs at 1 hour after psilocybin administration, with 1471, 1813, 2371, and 2492 DEGs detected in L6 IT, L5 IT, L4/5 IT, and L2/3 IT neurons, respectively (FDR < 0.05; **Fig. 3c**). At this early time point, psilocybin caused both increases and decreases in gene expression, with 58-61% of DEGs upregulated and 39-42% downregulated across the IT cell types. The transcriptional response then waned precipitously, with an average of only 82, 117, and 3 DEGs detected at 2, 4, and 24 hours after psilocybin administration. Unexpectedly, the number of DEGs increased again at 72 hours, climbing back to exceed 1,000 genes in each of the four IT cell types. This time course of transcriptional response, characterized by early and late waves of gene expression changes, is a recurring theme across multiple cell types following psilocybin administration.

**Fig. 3:**
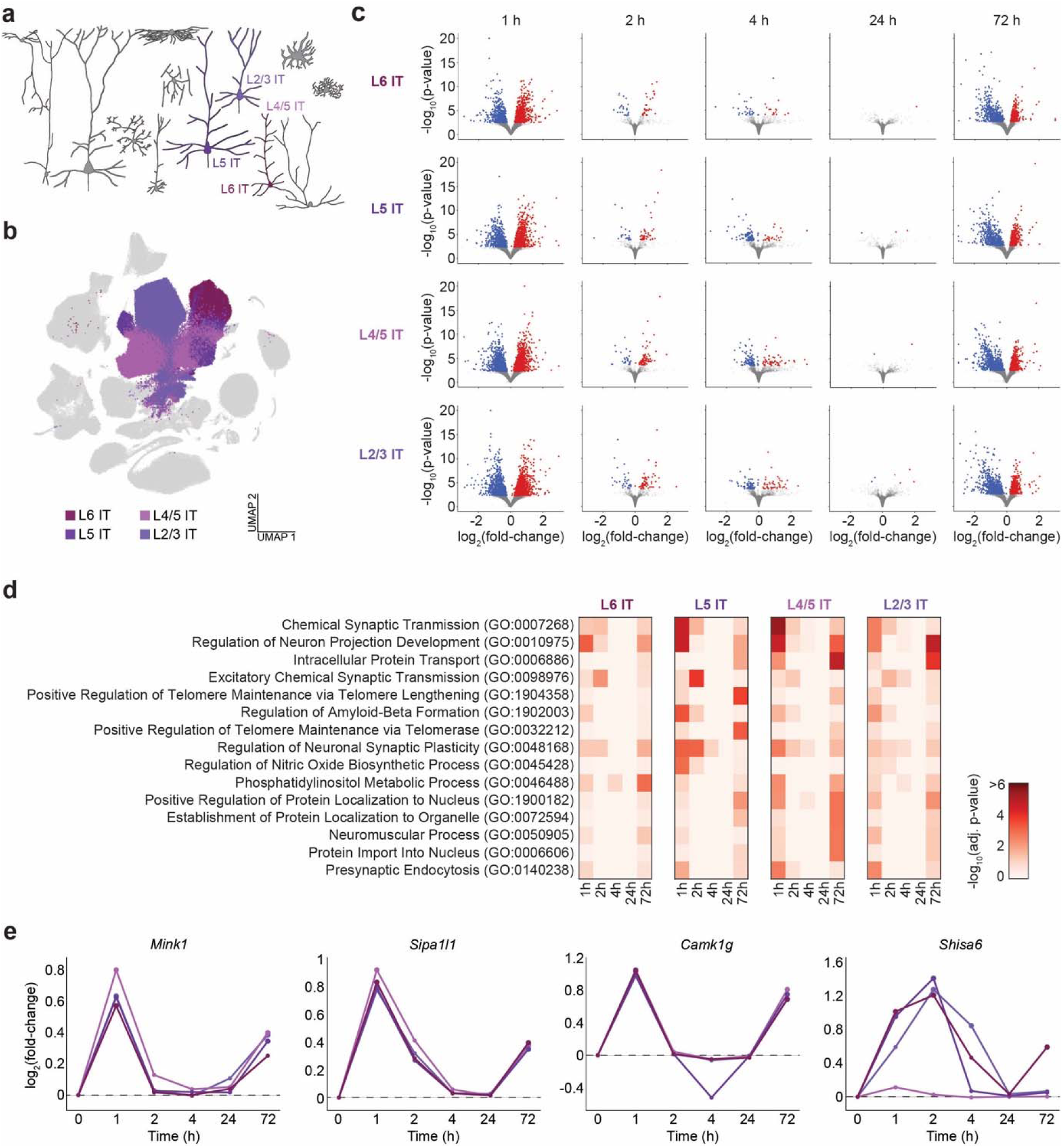
Psilocybin-induced transcriptional changes in frontal cortical IT excitatory neurons. **(a)** Schematic of the cortical microcircuit highlighting the four IT subtypes of excitatory neurons (L6 IT, L5 IT, L4/5 IT, and L2/3 IT). **(b)** UMAP representation of the snRNA-seq dataset highlighting the four cell types. **(c)** Differentially expressed genes (DEGs) identified by pseudobulk analysis for each time point for the four cell types. Each circle represents an individual gene. Red, upregulated with FDR < 0.05. Blue, downregulated with FDR < 0.05. **(d)** Top 15 terms from gene ontology enrichment analysis based on the upregulated DEGs, ranked based on the mean adjusted p-values across time points in the four cell types. **(e)** Expression levels of specific transcripts in the four cell types following psilocybin administration, normalized to controls. Color of the line denotes the cell type.

To gain insights into the biological functions associated with the psilocybin-induced transcriptional changes, we performed gene ontology (GO) enrichment analysis on the upregulated DEGs identified in IT neurons across time points. **Fig. 3d** shows the top 15 overrepresented GO terms when we considered the prevalence of each term averaged across all time points and for all four cell types. The top categories are processes related to synaptic plasticity and excitatory neurotransmission, including chemical synaptic transmission, regulation of neuron projection development, excitatory chemical synaptic transmission, and regulation of neuronal synaptic plasticity. Also notable are pathways related to protein synthesis and intracellular trafficking, such as intracellular protein transport, positive regulation of protein localization to nucleus, establishment of protein localization to organelle, and protein import into nucleus. To illustrate the genes contributing to these enriched pathways, we visualized select upregulated genes including *Mink1*, *Sipa1l1*, *Camk1g*, and *Shisa6* (**Fig. 3e**). *Mink1* encodes a kinase that maintains dendrite complexity by regulating actin cytoskeletal dynamics and AMPA receptor trafficking^45^. *Sipa1l1* is transcript for the SPAR1 protein, which localizes to the postsynaptic density and complexes with PSD-95 and NMDA receptors and contributes to spine head enlargement^46^. *Camk1g* encodes a component of a calcium- and calmodulin-dependent protein kinase complex involved in dendritic growth^47^. Select genes were upregulated over more extended durations after psilocybin, such as *Shisa6*, which contributes to anchoring AMPA receptors into postsynaptic compartments in dendrites^48^. We note that users of the web portal can generate similar expression plots for any genes of interest.

We next examine DEGs in the other excitatory neuron subtypes, including L5 ET, L6b, L6 CT, and L5 NP neurons **(Fig. 4a, b).** These excitatory cell populations also exhibit a biphasic pattern of transcriptional changes, with an initial peak at 1 hour and then late response at 72 hours after psilocybin administration (**Fig. 4c**). Gene ontology analysis revealed similar biological themes, including the same top enriched pathway of chemical synaptic transmission, as well as related yet distinct processes such as glutamatergic synaptic transmission and glutamate receptor signaling pathway (**Fig. 4d**). We plotted the expression profiles for several upregulated genes (**Fig. 4e**), such as *Nos1ap*, which binds neuronal nitric oxide synthase and plays a role in dendrite growth and NMDA receptor-dependent responses^49^. There are also genes with more extended responses including *Grin2a* and *Gria1*, which encode the NR2A subunit of NMDA receptor and GluA1 subunit of AMPA receptor, respectively, with clear roles in mediating glutamatergic transmission and excitatory synaptic plasticity^50, 51^. *Sorcs3* encodes a neuronal receptor that regulates intracellular protein transport^52^ and has also been implicated in synaptic plasticity^53^.

**Fig. 4:**
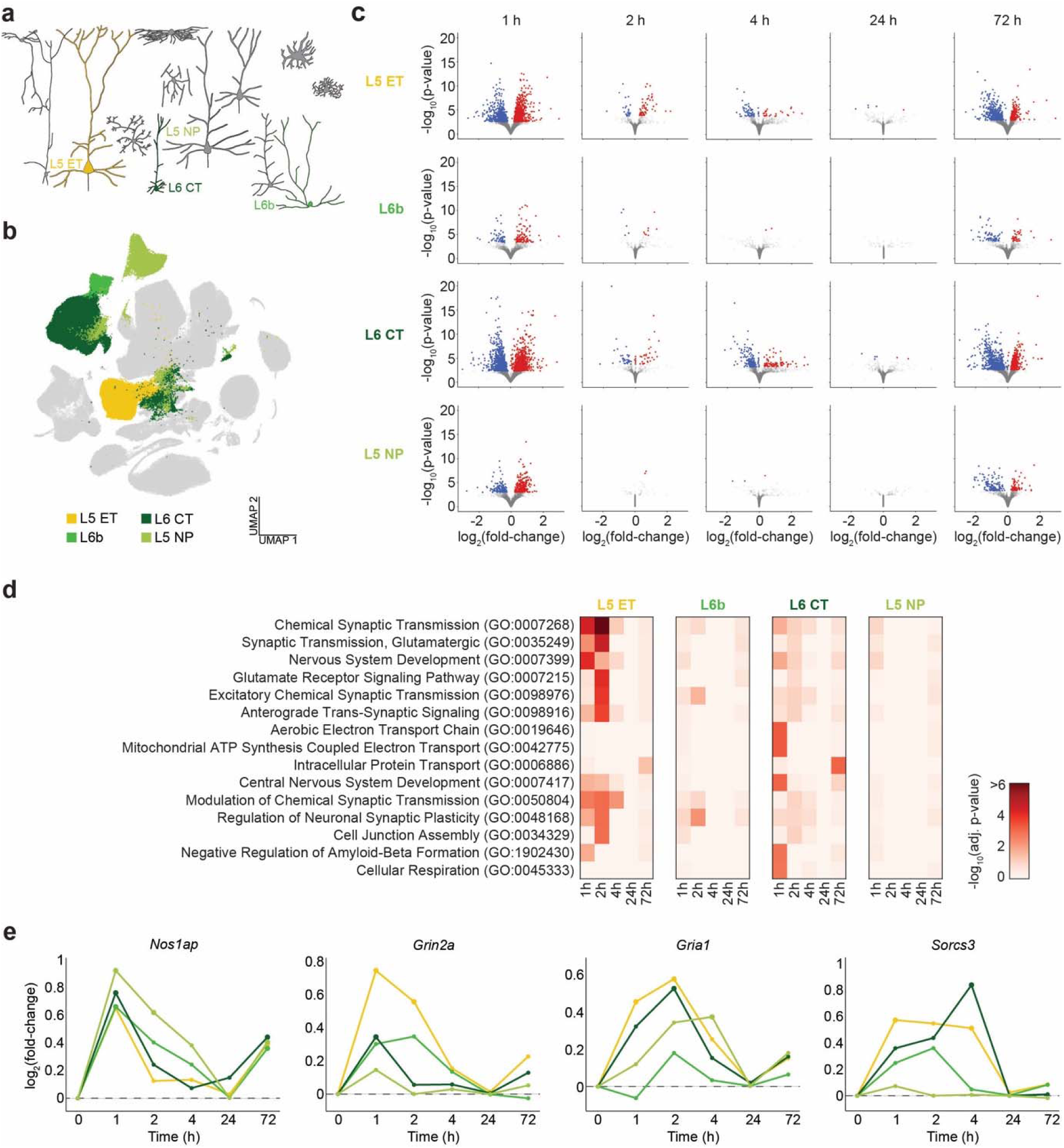
Psilocybin-induced transcriptional changes in other excitatory neuronal subtypes. **(a)** Schematic of the cortical microcircuit highlighting the four non-IT subtypes of excitatory neurons (L5 ET, L6b, L6 CT, L5 NP). **(b)** UMAP representation of the snRNA-seq dataset highlighting the four cell types. **(c)** Differentially expressed genes (DEGs) identified by pseudobulk analysis for each time point for the four cell types. Each circle represents an individual gene. Red, upregulated with FDR < 0.05. Blue, downregulated with FDR < 0.05. **(d)** Top 15 terms from gene ontology enrichment analysis based on the upregulated DEGs, ranked based on the mean adjusted p-values across time points in the four cell types. **(e)** Expression levels of specific transcripts in the four cell types following psilocybin administration, normalized to controls. Color of the line denotes the cell type.

### Psilocybin-induced transcriptional changes in frontal cortical GABAergic neurons contribute to mitochondrial function and metabolic processes

For GABAergic neurons, we characterized the transcriptional responses in Vip, Lamp5, Pvalb, and Sst cell types (**Fig. 5a, b**). The magnitude of the response varied across interneuron subtypes. At 1 hour after psilocybin administration, robust responses were detected in Pvalb and Sst interneurons, with 933 and 662 DEGs respectively, but changes in Vip and Lamp5 interneurons were comparatively modest, with 154 and 80 DEGs (**Fig. 5c**). Unlike excitatory neurons, GO enrichment analysis revealed that DEGs in GABAergic neurons had enriched pathways related to mitochondrial function and metabolic processes, such as aerobic electron transport chain, mitochondrial ATP synthesis coupled electron transport, and canonical glycolysis (**Fig. 5d**). We plotted select genes associated with the top pathways, including *Cox6c*, *Ndufs2*, *Kcnq3*, and *Amigo1* (**Fig. 5e**). *Cox6c* and *Ndufs2* encode key components of cytochrome c oxidase in the mitochondrial electron transport chain^54^ and mitochondrial complex I^55^, respectively. *Kcnq3* and *Amigo1* encode the subunits of a voltage-gated potassium channel to regulate neuronal excitability. Particularly abundant in Pvalb interneurons, *Kcnq3* and *Amigo1* were highlighted in the top enriched process of monoatomic cation transmembrane transport, possibly linking the use of metabolic energy to regulate cellular excitability. Overall, the analyses indicate that psilocybin engages transcriptional programs in frontal cortical GABAergic neurons that are associated with mitochondrial function and metabolic processes, distinguishing their response from the synaptic plasticity-related programs observed in excitatory neurons.

**Fig. 5:**
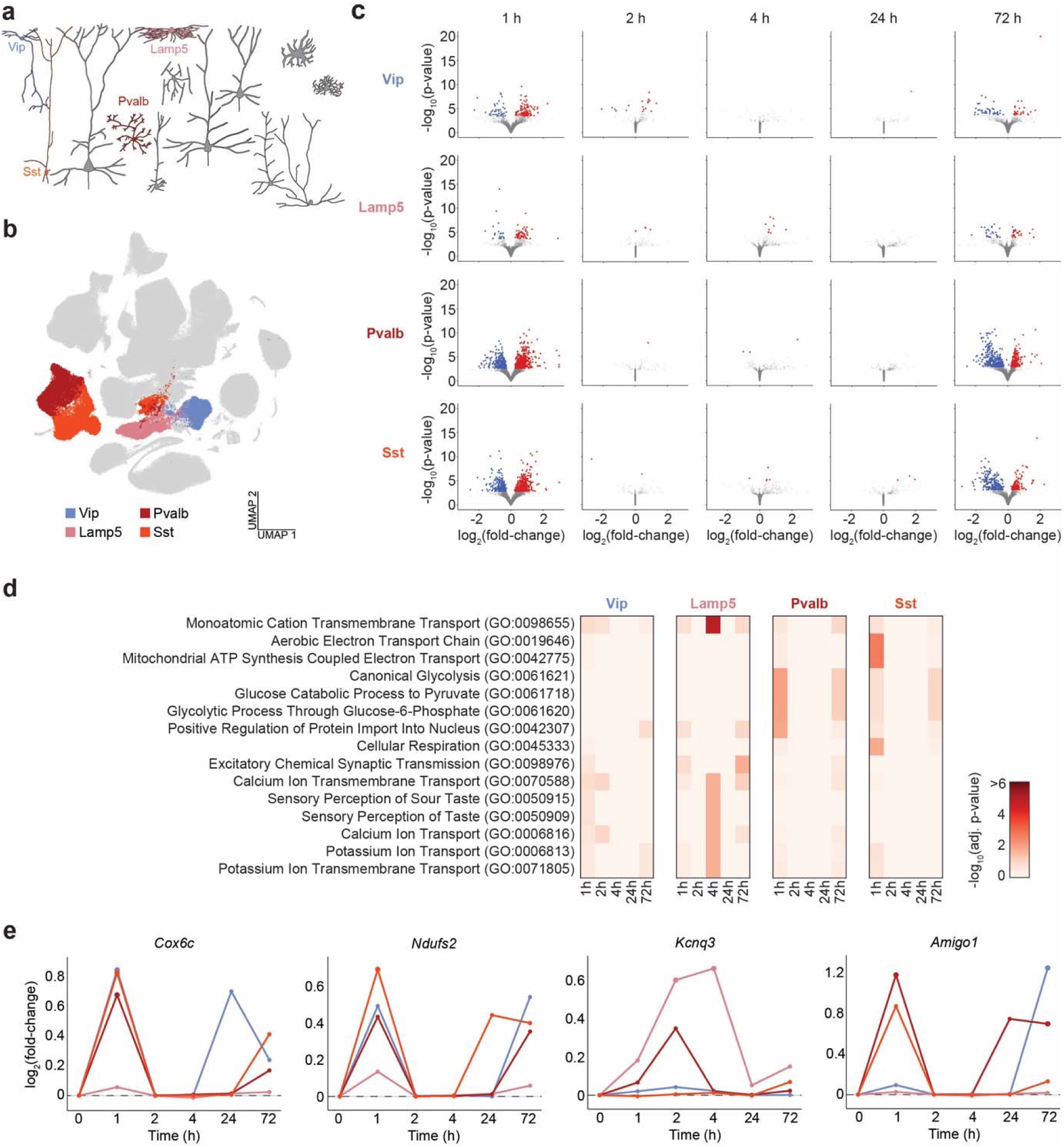
Psilocybin-induced transcriptional changes in frontal cortical GABAergic neurons contribute to mitochondrial function and metabolic processes. **(a)** Schematic of the cortical microcircuit highlighting the four major classes of GABAergic neurons (Vip, Lamp5, Pvalb, Sst). **(b)** UMAP representation of the snRNA-seq dataset highlighting the four cell types. **(c)** Differentially expressed genes (DEGs) identified by pseudobulk analysis for each time point for the four cell types. Each circle represents an individual gene. Red, upregulated with FDR < 0.05. Blue, downregulated with FDR < 0.05. **(d)** Top 15 terms from gene ontology enrichment analysis based on the upregulated DEGs, ranked based on the mean adjusted p-values across time points in the four cell types. **(e)** Expression levels of specific transcripts in the four cell types following psilocybin administration, normalized to controls. Color of the line denotes the cell type.

### Psilocybin-induced transcriptional changes in non-neuronal cells

Our analysis of control samples showed negligible levels of serotonin receptor transcripts in non-neuronal cells in the mouse medial frontal cortex (**Fig. 2**), yet surprisingly we found that they displayed transcriptional responses to psilocybin. The most responsive non-neuron populations were astrocytes and oligodendrocytes, with 776 and 789 DEGs at 1 hour after administration respectively (**Fig. 6a, b**). There were fewer but still notable changes involving 202 and 64 DEGs for oligodendrocyte precursor cells and microglia. GO analysis revealed pathways related to neuronal connectivity, such as neuron projection guidance, axon guidance, axonogenesis, and chemical synaptic transmission (**Fig. 6d**). As exemplars, we plotted the time courses for *Nlgn3*, *Lrp1*, *Aldoc*, and *Lars2* (**Fig. 6e**). *Nlgn3* was upregulated by psilocybin particularly in oligodendrocytes and oligodendrocyte precursor cells, where it plays a crucial role in their differentiation and myelination^56, 57^. *Lrp1* encodes a multicargo transporter that interacts with insulin-like growth factor to promote survival and proliferation, including in astrocytes^58^. *Aldoc* is the transcript for aldolase, which acts as a checkpoint for proliferation of oligodendrocyte precursor cells^59^. *Lars2* contributes to mitochondrial regulation and metabolic homeostasis and, interestingly, shows notable response preferentially in the late phase at 72 h. The results demonstrate that psilocybin induces transcriptional responses in non-neuronal cells in the medial frontal cortex, likely indirectly via circuit-level mechanisms.

**Fig. 6:**
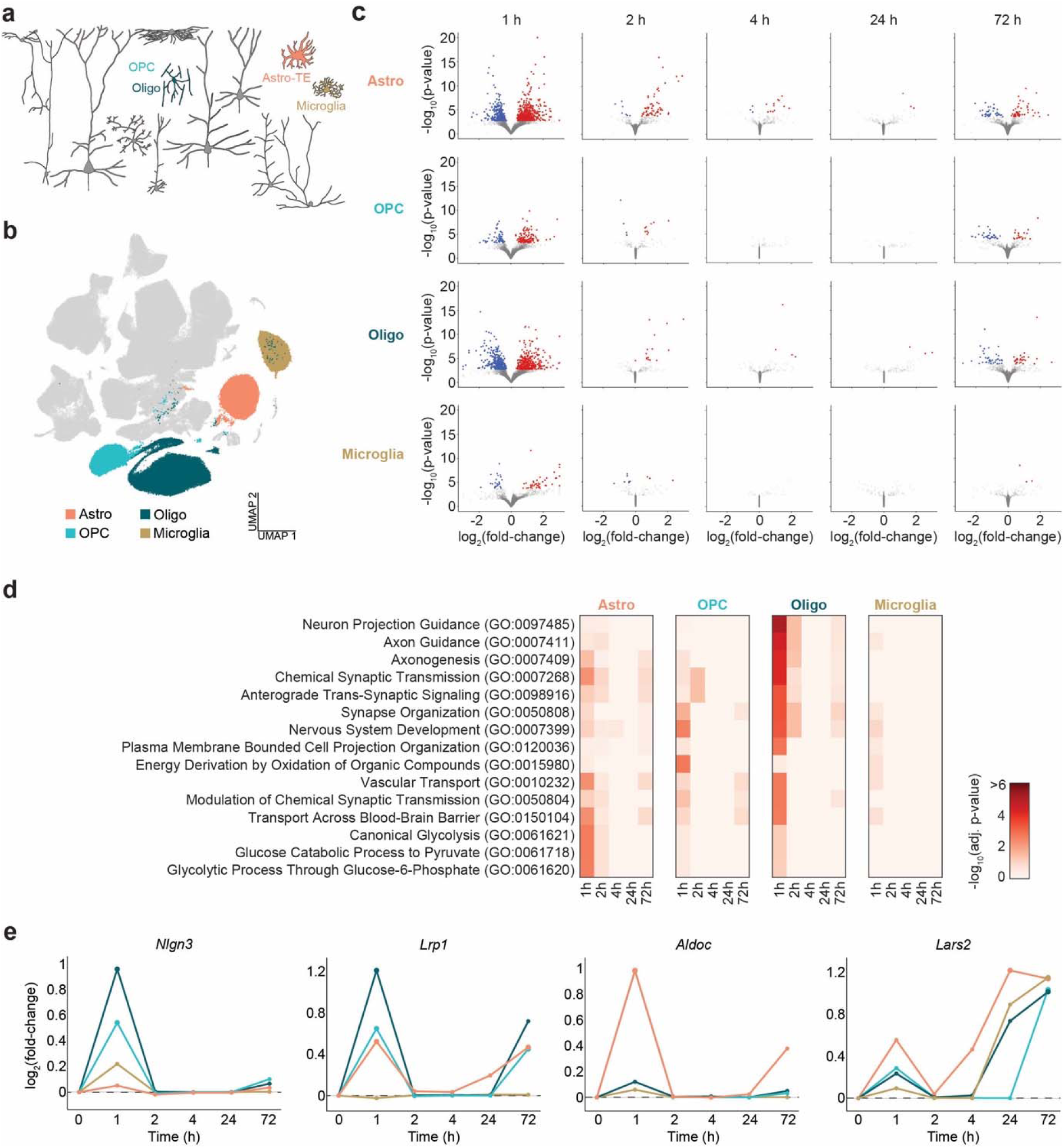
Psilocybin-induced transcriptional changes in non-neuronal cells. **(a)** Schematic of the cortical microcircuit highlighting the four non-neuronal cell types (Astro, OPC, Oligo, Microglia). **(b)** UMAP representation of the snRNA-seq dataset highlighting the four cell types. **(c)** Differentially expressed genes (DEGs) identified by pseudobulk analysis for each time point for the four cell types. Each circle represents an individual gene. Red, upregulated with FDR < 0.05. Blue, downregulated with FDR < 0.05. **(d)** Top 15 terms from gene ontology enrichment analysis based on the upregulated DEGs, ranked based on the mean adjusted p-values across time points in the four cell types. **(e)** Expression levels of specific transcripts in the four cell types following psilocybin administration, normalized to controls. Color of the line denotes the cell type.

### Comparing the time-dependent and cell type-specific transcriptional responses between psilocybin and ketamine

To facilitate comparisons across cell types and time points, we summarized the number of upregulated and downregulated DEGs following psilocybin administration (**Fig. 7a**). Psilocybin-induced transcriptional responses exhibited a biphasic temporal profile, consisting of an early wave of gene expression at 1 hour followed by a late response at 72 hours after drug administration. This visualization also highlights excitatory neurons as the cell class showing the highest number of DEGs, relative to the more modest number of DEGs in select GABAergic and non-neuronal cell types. Interestingly, when we performed a similar analysis for ketamine-treated animals, a different time course emerged. In contrast to psilocybin, ketamine induced fewer DEGs, which peaked at around 2 and 4 hours after administration (**Fig. 7b**).

**Fig. 7:**
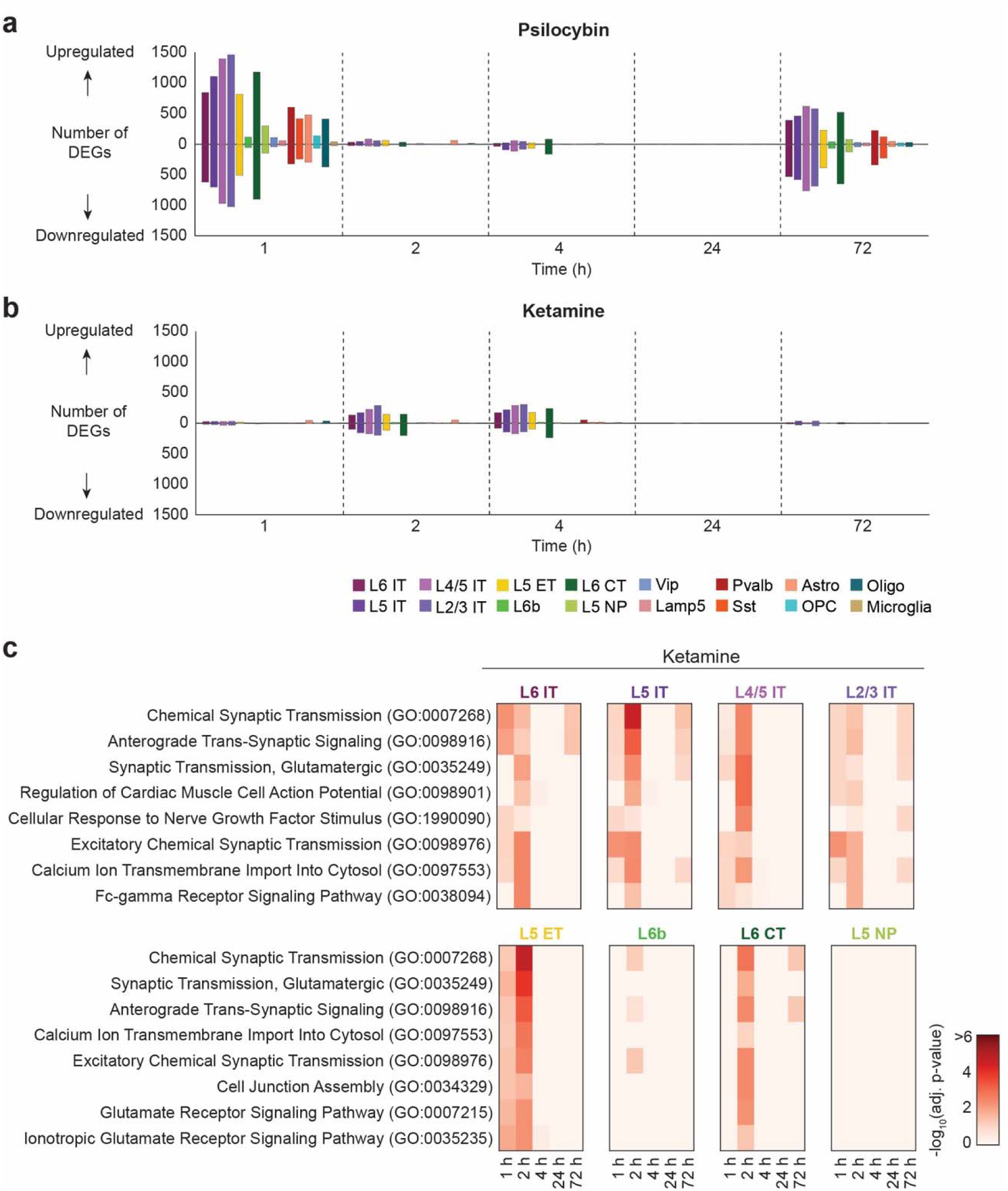
Distinct time courses of transcriptional responses evoked by psilocybin and ketamine. **(a)** The number of upregulated and downregulated DEGs (FDR < 0.05) identified by pseudobulk analysis for the 16 cell types at each time point following psilocybin administration. The color of the bar corresponds to the cell type, using the same color scheme as Fig. 1b. **(b)** Similar to (a) for ketamine. **(c)** Top 8 terms from gene ontology enrichment analysis based on the upregulated DEGs following ketamine administration, ranked based on the mean adjusted p-values across time points in the four cell types for IT neurons (top row) and non-IT excitatory neurons (bottom row).

For completeness, we performed the differential expression analyses including gene ontology enrichment analyses for ketamine-treated animals as well, for all 16 annotated cell types including the IT excitatory neuron subtypes (**Supplementary Fig. 4**), other excitatory neuron subtypes (**Supplementary Fig. 5**), inhibitory neuron subtypes (**Supplementary Fig. 6**), and non-neuronal cell types (**Supplementary Fig. 7**). Despite the different temporal dynamics, we observed examples of convergent biological pathways engaged by both ketamine and psilocybin. For instance, in excitatory neuron subtypes, the top enriched GO terms for the ketamine samples were chemical synaptic transmission, anterograde trans-synaptic signaling, and glutamatergic synaptic transmission (**Fig. 7c**). These findings indicate that both compounds, despite acting through distinct receptors, induce transcriptional programs associated with synaptic plasticity and excitatory neurotransmission.

## DISCUSSION

The present study applied single-nucleus transcriptomics to determine how psilocybin and ketamine alter gene expression in the mouse medial frontal cortex. As a resource, we annotated the dataset according to 8 major excitatory neuron types, 4 GABAergic cell types, and 4 non-neuronal cell types. We developed an interactive web portal (https://psilo-seq.kwanlab.org/) to facilitate data exploration. As examples of the insights enabled by this resource, we show that excitatory neurons, GABAergic neurons, and non-neuronal cells exhibit differentially expressed genes associated with distinct biological processes. We also find that psilocybin’s impact is time-dependent with early and late phases of transcriptional responses, which was distinct from ketamine’s time course. Together, the dataset provides a resource that unveils the dynamics of transcriptional responses in frontal cortical cell types after a single dose of psilocybin or ketamine.

The bolus injection of psilocybin causes rapid elevation of drug concentration in the central nervous system followed by full clearance within a few hours. This pharmacokinetic profile hints at a temporal pattern of transcriptional changes, and here we identified two distinct phases: an early phase at 1 – 2 hours followed by a late phase at 72 hours, each marked by a substantial number of differentially expressed genes. Indeed, a couple of studies have reported long-lasting changes in gene expression after the administration of psychedelics for one or few weeks^60, 61^. Our findings echo the framework that there are often multiple waves of transcription following a perturbation^11, 62^. For example, there are early and late response genes in visual cortical cells in dark-reared animals after their first experience of visual stimuli^63^. It is also possible that the long-term changes in gene expression reflect effects of psychedelics on alternative splicing or epigenomic modifications^60, 64^. It is tempting to speculate that the multiple phases of transcriptional responses observed in this study may reflect therapeutic mechanisms for ketamine and psilocybin at different time scales.

Notwithstanding the new insights into the cell type specificity and timing of drug-evoked transcriptional responses, several of the differentially expressed genes and pathways identified here are consistent with prior studies. Regulation of transcripts associated with synaptic plasticity and glutamatergic transmission is expected given previous reports of psilocybin- and ketamine-evoked structural plasticity in the medial frontal cortex^6–10^. Notably, a pioneering study examining the effects of LSD on rat frontal cortex, which profiled about 3,000 genes, reported preferential regulation of genes involved in synaptic function, glutamatergic signaling, and cytoskeletal plasticity^19^. Mitochondrial biogenesis has also been linked to serotonergic signaling, particularly through the activation of 5-HT_2A_ receptors^65^. It is worth noting that multiple studies have reported increased expression of immediate early genes such as *Fos* and *Arc* following psychedelic administration, based on transcript quantifications^15, 19, 21^ or whole-brain immunohistochemistry^34, 66^. While we examined *Fos, Jun, Npas4, Arc, Egr1,* and *Egr2* in our dataset, only *Arc* and *Egr1* in astrocytes showed significant upregulation for psilocybin-treated animals at the 1-hour time point (**Supplementary Fig. 8a**) and none of these genes reached significance for ketamine at 1 hour (**Supplementary Fig. 8b**). The discrepancy could be due to methodological differences: our approach measures nuclear RNA, which exhibits more rapid and transient dynamics than total cellular RNA or protein. Previous work comparing activity-dependent transcriptional changes across measurement techniques found that such changes are more difficult to detect using snRNA-seq^67^.

A weakness of this study is that the psilocybin- and ketamine-treated groups were compared against controls that did not receive an intraperitoneal injection. The injection itself can induce transcriptional responses, particularly in stress-sensitive cell types and brain regions. Ideally, the study would have included vehicle-injected controls collected at each corresponding time point (1, 2, 4, 24, and 72 hours). However, the large number of additional samples required for such a design was prohibitive given the costs. To mitigate this weakness, we increased the size of the control cohort, anticipating the importance of having a robust reference dataset for differential expression analyses. In addition, head-to-head comparison of the psilocybin and ketamine datasets revealed markedly different temporal patterns of transcriptional responses (**Fig. 7a**). If the injection procedure were a major driver of the observed gene expression changes, we would expect the two datasets to exhibit similar temporal profiles. Therefore, any contribution of the injection protocol is likely modest relative to the drug-specific transcriptional effects.

There are limitations for this study. Transcriptomics is a rapidly growing field with numerous approaches for sequencing and analysis. Here, we chose to work with single nuclei instead of single cells, because we were concerned that cortical neurons with extended axons and dendrites may be damaged in protocols needed to dissociate single cells, which can lead to aberrant readout of gene expression^67, 68^. Sequencing with single nuclei may better reflect recent changes in *de novo* transcription, and has been shown to yield excellent sensitivity and reliable classification of cell types^69^. Moreover, previous studies have identified sex as an important biological variable that can influence the behavioral effects of psychedelics^70–72^. However, the current dataset was not sufficiently powered to support a rigorous analysis of sex-specific transcriptional response.

In summary, this dataset serves as a resource for investigating the transcriptional responses to psilocybin and ketamine across frontal cortical cell types. We anticipate the growing number of studies revealing the molecular and cellular impact of psilocybin and ketamine will eventually reveal mechanisms of action that can accelerate the development of more effective and safer drugs for treating mental illnesses.

## METHODS

### Animals

Animal care and experimental procedures were approved by the Animal Care & Use Committee (IACUC) at Yale University and Cornell University. C57BL/6J (Stock No. 000664) mice were purchased from Jackson Laboratory and habituated in our animal facility at Yale University for 1 week. Animals were housed in same-sex groups of 2-5 mice per cage in a temperature-controlled room, operating in normal 12 hr light - 12 hr dark cycle (8:00 AM to 8:00 PM for light). Food and water were available ad libitum. Animals were randomly assigned to different experimental groups. 8-week-old animals were injected with psilocybin (1 mg/kg, i.p, prepared from working solution, which was made fresh monthly from powder; Usona Institute) or ketamine (10 mg/kg, i.p; Zoetis), returned to their home cage, and used 1, 2, 4, 24, or 72 hours after injection. Additional age-matched mice were used with no drug administration as controls. More details for the animals, including their sex and assignment to each treatment group, are included in **Supplementary Table 1**.

### Tissue collection and generation of single-cell suspensions

For tissue collection, decapitation immediately followed cervical dislocation and brains were immediately dissected from the skull. The dorsal medial frontal cortex, consisting of ACAd and medial MOs, was microdissected in cold RNAse-free 1x PBS, flash frozen in ethanol and dry ice, and stored at -80°C. Samples were shipped overnight on dry ice to the University of Michigan for processing. Nuclei were isolated and processed for single-nucleus sequencing using RNase-free materials and reagents. Fresh buffers were made immediately before nuclei processing. Buffers were made in accordance with recommended protocol (CG000393 Rev A, “Nuclei isolation from adult mouse brain tissue for single cell RNA sequencing”, 10x Genomics). Lysis buffer consisted of 10 mM Tris-HCl ph7.4 (Millipore-Sigma T2194), 10 mM NaCl (Millipore-Sigma 59222C), 3 mM MgCl_2_ (Millipore-Sigma, M1028), 0.1% Nonidet P40 (Millipore-Sigma 74385), and nuclease-free water (Invitrogen AM9932). Wash buffer consisted of 1% BSA solution (Thermo Fisher Scientific AM2616), 0.2U/µl RNase Inhibitor (Millipore-Sigma 3335399001), and 1x PBS (Gibco, 10010023). Tissue was dounce homogenized in 1 mL lysis buffer and centrifuged at 500g for 5 minutes at 4°C. Supernatant was removed and samples were resuspended in 1mL of wash buffer. Samples were strained through 40 µm cell strainers (Bel-Art H13680-0040) and centrifuged again at 500 g for 5 minutes at 4°C. Supernatant was removed and samples were resuspended in 1mL of wash buffer and passed through 40 µm cell strainers once more. Nuclei were visually checked with Trypan Blue Stain (Thermo Fisher Scientific T10282) to ensure minimal debris and efficient lysis. Nuclei were stained with 0.25 µL 7AAD (Invitrogen A310) and immediately flow sorted and counted on a MoFlo Astrios (Beckman Coulter) at the North Campus Research Complex Flow Core at University of Michigan to reduce ambient RNA. Samples were centrifuged and resuspended in wash buffer at a concentration of 1,000 cells/µL.

### Single cell library preparation and sequencing

Library prep and next-generation sequencing were carried out with the 10x Genomics Chromium Single Cell 3’ v3 platform in the Advanced Genomics Core at the University of Michigan. Resulting libraries were sequenced on the Illumina NovaSeq S4 300 cycle, at a target depth of 25,000 reads per cell and a target of 10,000 cells per sample.

### Single cell quality control, clustering, and cell type identification

Single nucleus RNA sequencing at the University of Michigan Advanced Genomics Core returned demultiplexed FASTQ files from all 49 samples. The 10x Genomics Mouse mm10 (GENCODE vM23/Ensembl98) reference version 2020-A was downloaded as a reference genome. FASTQ files were processed with 10x Genomics cellranger count^73^ version 7.1.0 and aligned to this reference, including intronic reads. Preprocessing and further analysis relied on anndata^74^ (version 0.12.7) and scanpy^75^ (version 1.11.5). Filtered feature-barcode matrices for all samples and all drug treatments were imported to AnnData objects and concatenated using anndata.concat() on the intersection of genes.

Out of the 49 samples, 7 were excluded to yield 42 samples for final analysis. Sample IDs 39, 40, 41, and 42 were excluded for oversequencing, because we incorrectly ordered for a targeted depth of 250,000 reads per nucleus at the facility. This introduced undesirable technical effects that could not be removed despite subsetting and batch integration attempts. Sample ID 290 was excluded due to low cell number, containing only 968 cells after sequencing compared to the median of 8,216. Sample ID 299 was excluded because it had the second lowest cell count at 3,372, was flagged for a high fraction of reads mapping antisense to genes by the cellranger pipeline, and analysis showed little transcriptional separation among cells. Sample 301 was excluded because its nuclei formed distinct tails extending from robust clusters in the UMAP embedding. Although the exact cause was unclear, this could be due to low-quality nuclei with reduced RNA content.

Analysis with the cellranger pipeline confirmed a mean depth of 29,562 reads per nucleus across all samples, consistent with the targeted read depth. Most samples exceeded the target depth of 25,000 reads per nucleus. Nine samples had sequencing depths below 25,000 reads per nucleus, with the lowest being 20,694 reads per nucleus. Thus, all samples exceeded the threshold of 20,000 reads per nucleus recommended by 10x Genomics. We also examined sequencing saturation, which ranged from 35–65% across samples.

Then we proceeded to process the data through several standard steps in cellranger. First, only reads that map confidently in the sense orientation to a single gene are retained. Second, for deduplication, reads sharing the same cell barcode and unique molecular identifier (UMI) are collapsed into a single transcript count because they originate from PCR amplification of the same RNA molecule. psilocybin-treated samples had 8482±1086 reads per nucleus (mean ± standard deviation), ketamine-treated samples had 8812±1436 reads per nucleus, and control samples had 8343±602 reads per nucleus.

Next, we performed additional filtering for quality control. All genes detected in fewer than 5 unique nuclei or having fewer than 5 total transcript counts were removed. All nuclei with fewer than 3 unique genes detected or having fewer than 3 total transcript counts were removed. Multiplet nuclei were identified and removed with scanpy.pp.scrublet including a batch key identifying individual samples. 13 samples were identified as homogenous, where scrublet identified few or no multiplets. To perform multiplet removal in these samples, the largest multiplet score threshold from the non-homogenous samples was identified and applied to the homogeneous samples. Nuclei with low transcript counts on a per-sample basis (<1000 counts and less than 3 mean absolute deviations below the sample median) were removed. Cells with high mitochondrial or ribosomal counts on a per-sample basis (>20% of counts or more than 3 mean absolute deviations above the sample median) were removed. Ribosomal genes were based on the Ribosomal Protein Gene Database^76^ (http://ribosome.med.miyazaki-u.ac.jp/). The ubiquitously and highly expressed *Malat1* gene was removed from analysis. At the end, psilocybin-treated samples contained 8,049±1,115 usable reads per nucleus (mean ± standard deviation), ketamine-treated samples contained 8,414±1,588 usable reads per nucleus, and control samples contained 7,689±826 usable reads per nucleus.

Each cell’s reads were normalized to counts per million (CPM) with scanpy.pp.normalize_total, and a natural logarithm was taken with scanpy.pp.log1p^77^. This yielded ln(CPM+1), which was used for all downstream expression analysis except pseudobulk. We identified the top 5000 highly variable genes using scanpy.pp.highly_variable_genes with flavor=‘seurat’. Batch corrected embedding was performed with scVI^78^ version 1.4.1 on these highly variable genes from raw RNA counts. The embedding model used default settings, having 10 latent dimensions, one 128-dimensional hidden layer in both the encoder and decoder, 10% dropout rate, and zero-inflated negative binomial likelihood. Nearest neighbors and UMAP was run on the learned batch-corrected embeddings with scanpy default settings. Cell types were annotated using the Allen Brain Atlas MapMyCells tool^35^ version 1.6.1 from raw RNA counts using hierarchical mapping, with 10x Whole mouse brain taxonomy (CCN20230722) as reference. Annotations from the ‘Subtype’ level were used, specifically identifying 004 L6 IT CTX Glut, 005 L5 IT CTX Glut, 006 L4/5 IT CTX Glut, 007 L2/3 IT CTX Glut, 022 L5 ET CTX Glut, 029 L6b CTX Glut, 030 L6 CT CTX Glut, 032 L5 NP CTX Glut, 046 Vip Gaba, 047 Sncg Gaba, 049 Lamp5 Gaba, 052 Pvalb Gaba, 053 Sst Gaba, 319 Astro-TE NN, 326 OPC NN, 327 Oligo NN, 333 Endo NN, and 334 Microglia NN. There were fewer cells with annotations 047 Sncg Gaba and 333 Endo NN, and therefore they were excluded from further analysis. Cells with annotations not included in this list were marked as “Other” and also excluded from further analysis. Marker genes for the remaining cell types were verified with scanpy.tl.rank_genes_groups function and the Wilcoxon rank-sum test, and plotted with scanpy.pl.heatmap.

### Serotonin receptor transcript expression

To examine cell type-specific expression patterns of psilocybin’s 5-HT receptor targets, we extracted *Htr1a*, *Htr2a*, and *Htr2c* transcript levels from the control samples in our snRNA-seq dataset. Cell type-specific patterns were visualized on a per-cell basis on the dataset’s UMAP representation. Group-wise variation among distinct cell subtypes was visualized with violin plots. UMAPs were created using matplotlib and the scanpy UMAP embeddings of the dataset, while violin plots were constructed using scanpy.pl.violin(). The scale of expression was unified across all genes by identifying the maximum expression level.

Transcript levels from our snRNA-seq dataset were compared to publicly available single-cell sequencing dataset from the Allen Institute for Brain Science (https://portal.brain-map.org/atlases-and-data/rnaseq/mouse-whole-cortex-and-hippocampus-smart-seq). Briefly, this independent dataset used SMARTseq-v4 to analyze 76,381 single cell transcriptomes across the mouse cortex and hippocampus, and cluster these cells into transcriptional cell subtypes^38, 79^. We selected the 16,953 cells sampled in frontal cortex regions (ACA, ALM, ORB, and PL-ILA), including 640 L6 IT, 1541 L5 IT, 3118 L4/5 IT, 1403 L2/3 IT, 471 L5 PT, 579 L6b, 2159 L6 CT, 1009 L5/6 NP, 1708 Vip, 1099 Lamp5, 1174 Pvalb, 1573 Sst, 277 Astro, 118 Oligo, and 84 Micro_PVM, as identified and annotated already by the Allen Institute. Transcript levels for 5-HT receptors (*Htr1a*, *Htr1b*, *Htr1d*, *Htr2a*, *Htr2b*, *Htr2c*, *Htr3a*, *Htr3b*, *Htr4*, *Htr5a*, *Htr5b*, *Htr6*, *Htr7*), as well as cell type markers for excitatory (*Slc17a7*) or inhibitory (*Gad1*, *Vip*, *Ndnf*, *Pvalb*, *Sst*) neurons were extracted for each cell subtype. Expression levels were depicted in a dot plot showing log_2_(CPM+1) per cell type, with dot size scaled by the percentage of cells of that type with non-zero transcript reads. Processing and visualization of Allen Institute data were done using MATLAB scripts.

### Differential gene expression using pseudobulk

We performed cell type-specific differential expression analysis using pseudobulk techniques to identify genes of interest while minimizing the risk of false discoveries. For pseudobulk, raw transcript counts were sum aggregated within each sample-cell type combination using ADPBulk version 0.1.4. Differential expression testing was performed with pyDESeq2^80^ version 0.5.3 with fold change shrinkage. Genes with less than 10 total counts were excluded for the given comparison. Differential expression was measured with log_2_(fold-change) and p-values were adjusted false discovery rate (FDR) correction. Shrinkage was applied to the log_2_(fold-change) values with pyDEseq2. P-value correction was applied with the Benjamini-Hochberg procedure to control the false discovery rate. Genes were identified as significant with an FDR below 5%. plotnine version 0.15.2 was used to create volcano plots. Genes with log_2_(fold-change) or -log_1_0(p-value) beyond the bounds of the plot area were clamped to be displayed on the axis limits.

### Gene ontology analysis

Gene ontology (GO) analysis was performed using Enrichr^81–83^ and Gene Ontology Biological Processes 2026 database to identify biological processes that were overrepresented for genes that were differentially expressed. For each cell type at each time point, if there were ≥5 upregulated DEGs, then the list of upregulated DEGs was analyzed using Enrichr. It is important to perform this analysis against an appropriate set of background genes. If we used all genes in the genome, the results can be misleading because, for example, excitatory neurons already express many neuronal and synaptic genes at baseline, and they will show up as spuriously enriched if the background is set to all mouse genes. For this reason, we first determine the background genes for each cell type, by taking the output from DESeq2 and filtered the expressed genes, defined as having baseMean > 10 at any time point or drug condition. Compared to the total number of 26,100 genes tested in DESeq2, this step produced cell type-specific background lists with ∼8,000 – 15,000 genes (specifically, L6 IT: 12693 genes; L5 IT: 13108; L4/5 IT: 14951; L2/3 IT: 14557; L5 ET: 13190; L6b: 8233; L6 CT: 13721; L5 NP: 10272; Vip: 7937; Lamp5: 8923; Pvalb: 11404; Sst: 10629; Astro: 9308; OPC: 8250; Oligo: 9964; Microglia: 6898). The adjusted p-values were computed by Enrichr using the Benjamini-Hochberg procedure for multiple hypotheses testing correction. Results for each cell type were then considered in groups: IT neurons (L6 IT, L5 IT, L4/5 IT, L2/3 IT), non-IT excitatory neurons (L5 ET, L6b, L6 CT, L5 NP), inhibitory neurons (Vip, Lamp5, Pvalb, Sst), and non-neuronal cells (Astro, OPC, Oligo, Microglia). For each group, to select the top GO terms, we averaged the - log_10_(adjusted p-value) from GO analysis across the four cell types and across time points to rank the GO terms. To identify representative genes, we first determined the most frequent associated genes for each of the top 30 GO terms across all time points.

## DATA AVAILABILITY

The data can be access and downloaded via an interactive web portal (https://psilo-seq.kwanlab.org/about). Raw sequencing data associated with this study have been submitted to the NIH Sequence Read Archive under BioProject ID PRJNA1204073 (http://www.ncbi.nlm.nih.gov/bioproject/1204073). Processed data, including the AnnData h5ad file with processed and raw counts and the pseudobulk differential expression sqlite database, are available on Zenodo (https://doi.org/10.5281/zenodo.19666128).

## CODE AVAILABILITY

Computer code to process and analyze the snRNA-seq data is provided as Snakemake scripts, Jupyter notebooks, and R scripts on GitHub (https://github.com/thekwanlab/psilocybin_cortex_snRNAseq).

## ACKNOWLEDGEMENTS

We thank Rosemarie Terwilliger and Mandy Lam for coordination assistance and guidance on sample preparation. Psilocybin was generously provided by Usona Institute’s Investigational Drug & Material Supply Program; the Usona Institute IDMSP is supported by Alexander Sherwood, Robert Kargbo, and Kristi Kaylo in Madison, WI. This work was supported by NIH grants R01MH121848 (A.C.K.), R01MH128217 (A.C.K.), R01MH137047 (A.C.K.), R01NS125313 (K.Y.K.), R01MH126287 (K.Y.K.), One Mind – COMPASS Rising Star Award (A.C.K.), NIH training grant T32NS041228 (C.L.), NIH fellowships F30MH129085 (N.K.S.), and NCI P30CA046592 (Cancer Center Shared Resource: Single Cell and Spatial Analysis Shared Resource).

## CONTRIBUTIONS

C.L, K.Y.K., and A.C.K planned the study. C.L. and A.M.W. conducted the experiment. E.O. and Y.Q. performed the analysis. C.L. and A.M.W. assisted in the analysis. N.K.S. and A.C.K. analyzed the data from Allen Institute. M.J.G. contributed to the planning of the study. C.L., K.Y.K., and A.C.K. drafted the manuscript. All authors reviewed the manuscript before submission.

## DISCLOSURES

A.C.K. has been a scientific advisor or consultant for Boehringer Ingelheim, Eli Lilly, Empyrean Neuroscience, Freedom Biosciences, Helus Pharma, Otsuka, and Xylo Bio. A.C.K. has received research support from Intra-Cellular Therapies. The other authors report no financial relationships with commercial interests.

## SUPPLEMENTARY FIGURES

**Supplementary Fig. 1:** UMAP representations of the snRNA-seq data set.

**Supplementary Fig. 2:** Bootstrap probabilities as confidence metric for cell-type classification via MapMyCells

**Supplementary Fig. 3**: Unsupervised Leiden clustering versus MapMyCells

**Supplementary Fig. 4:** Ketamine-induced transcriptional changes in frontal cortical IT excitatory neurons

**Supplementary Fig. 5:** Ketamine-induced transcriptional changes in other excitatory neuronal subtypes

**Supplementary Fig. 6:** Ketamine-induced transcriptional changes in frontal cortical GABAergic neurons

**Supplementary Fig. 7:** Ketamine-induced transcriptional changes in non-neuronal cells

**Supplementary Fig. 8:** Immediate early genes

## SUPPLEMENTARY TABLES

**Supplementary Table 1.** Information regarding the samples.

**Supplementary Table 2.** The number of animals and cells in each experimental group.

**Supplementary Table 3.** The number of differentially expressed genes in each cell type and time point.

## SUPPLEMENTARY DATA

The lists of differentially expressed genes across cell types, time points, and drug treatment.

## Notes

http://www.ncbi.nlm.nih.gov/bioproject/1204073

https://github.com/thekwanlab/psilocybin_cortex_snRNAseq

https://psilo-seq.kwanlab.org/about

https://doi.org/10.5281/zenodo.19666128

### Summary of Updates

Revised manuscript and supplementary materials. Provided links to an interactive web portal and processed data.

