## Supplementary Figures for "Single-nucleus transcriptomics reveals cell type-specific and time-dependent effects of psilocybin and ketamine on gene expression"

**Supplementary Figures 1 - 8**

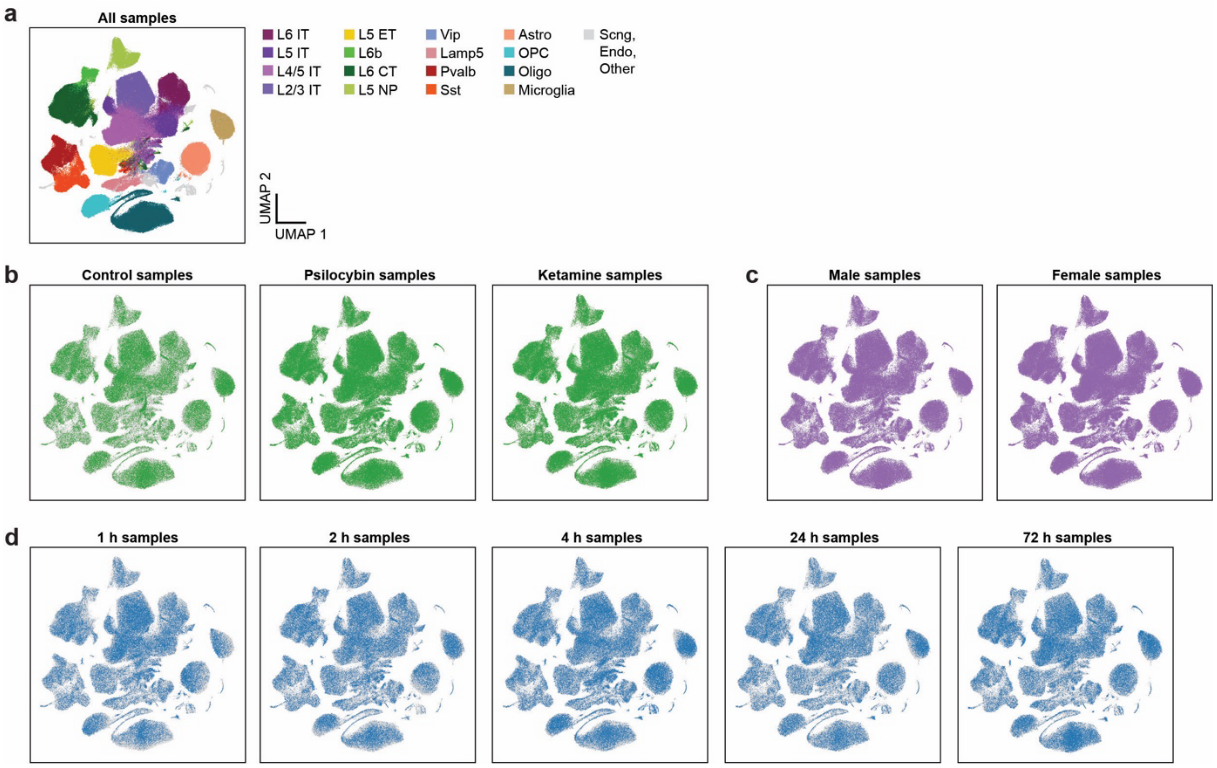


**Supplementary Fig. 1: UMAP representations of the snRNA-seq data set.**

**(a)** UMAP representation of the snRNA-seq dataset, including all samples. Color denotes the cell type.
**(b)** UMAP representations of the snRNA-seq dataset for control samples only, psilocybin samples only, or ketamine samples only. Colored circle, cell in the specified subset of samples. Gray circle, cell in all samples.
**(c)** UMAP representations of the snRNA-seq dataset for male samples only, or female samples only. Colored circle, cell in the specified subset of samples. Gray circle, cell in all samples.
**(d)** UMAP representations of the snRNA-seq dataset for samples only from 1 h time point, samples only from 2 h time point, samples only from 4 h time point, samples only from 24 h time point, or samples only from 72 h time point. Colored circle, cell in the specified subset of samples. Gray circle, cell in all samples.

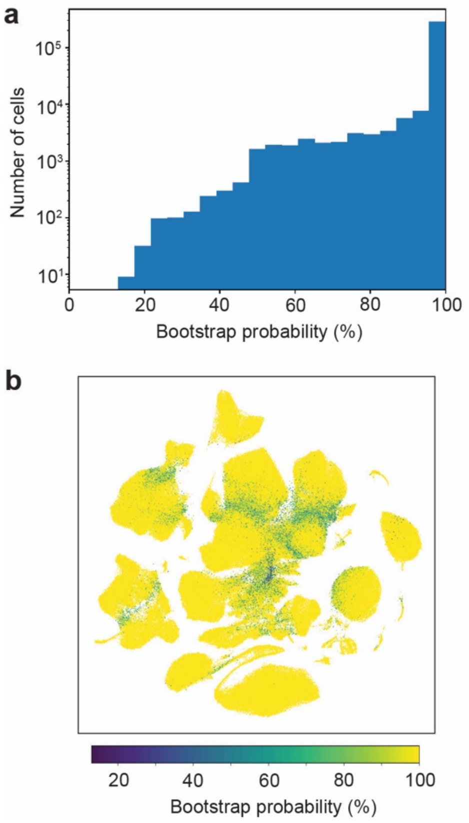


c

Proportion of cells with bootstrap probabiliy exceeding threshold (%)

| Threshold | 50% | 80% | 90% | 95% | 99% |
|---|---|---|---|---|---|
| 004 L6 IT CTX Glut | 98.6 | 91.0 | 87.3 | 83.9 | 75.3 |
| 005 L5 IT CTX Glut | 98.3 | 83.1 | 75.1 | 67.7 | 52.0 |
| 006 L4/5 IT CTX Glut | 99.0 | 90.0 | 85.9 | 82.4 | 74.6 |
| 007 L2/3 IT CTX Glut | 99.3 | 92.4 | 88.2 | 84.2 | 73.6 |
| 022 L5 ET CTX Glut | 99.6 | 97.9 | 96.6 | 95.4 | 93.0 |
| 029 L6b CTX Glut | 99.3 | 85.6 | 79.7 | 75.2 | 64.2 |
| 030 L6 CT CTX Glut | 99.9 | 96.4 | 93.4 | 89.5 | 74.1 |
| 032 L5 NP CTX Glut | 99.8 | 97.0 | 95.9 | 94.9 | 92.2 |
| 046 Vip Gaba | 99.9 | 98.2 | 97.2 | 96.0 | 91.6 |
| 047 Scng Gaba | 99.9 | 91.4 | 86.3 | 81.4 | 68.3 |
| 049 Lamp5 Gaba | 99.9 | 97.6 | 96.1 | 94.5 | 89.5 |
| 052 Pvalb Gaba | 100.0 | 98.1 | 96.6 | 95.0 | 90.0 |
| 053 Sst Gaba | 100.0 | 97.8 | 96.8 | 95.7 | 93.1 |
| 319 Astro-TE NN | 99.9 | 96.6 | 92.7 | 87.8 | 69.8 |
| 326 OPC NN | 100.0 | 99.3 | 99.3 | 99.0 | 98.2 |
| 327 Oligo NN | 100.0 | 99.8 | 99.7 | 99.6 | 99.5 |
| 333 Endo NN | 100.0 | 99.7 | 99.7 | 99.7 | 97.1 |
| 334 Microglia NN | 100.0 | 100.0 | 99.9 | 99.7 | 97.4 |
| Other | 94.0 | 86.3 | 84.1 | 82.4 | 77.4 |
| All cells | 99.4 | 94.4 | 91.5 | 88.7 | 80.7 |

**Supplementary Fig. 2: Bootstrap probabilities as confidence metric for cell-type classification via MapMyCells**
**(a)** The bootstrap probabilities for all cells in our dataset. MapMyCells provides bootstrap probability as a confidence metric for the annotation of each cell. Bootstrap probability is determined by iteratively applying the algorithm, and reporting the fraction of time the algorithm selects a particular cell type.
**(b)** UMAP representations of the snRNA-seq dataset. Color denotes the bootstrap probability.
**(c)** For each annotated cortical cell type, the percentage of cells in our dataset that have bootstrap probability above a certain threshold. Gray shading denotes the corresponding proportion value.

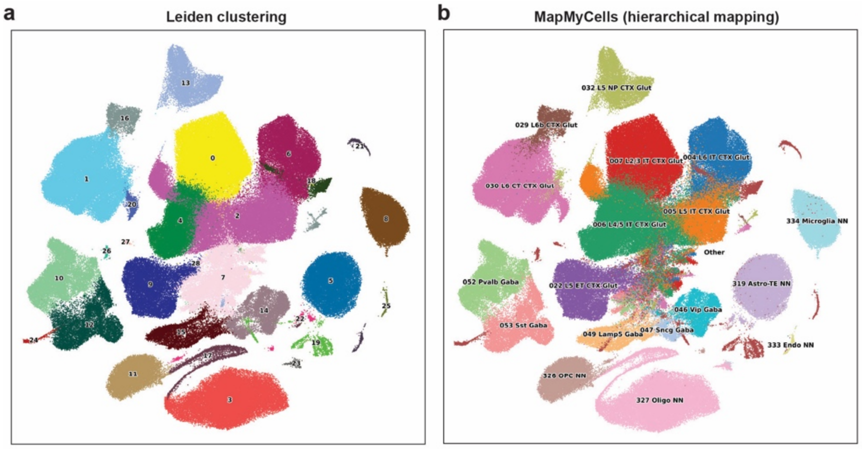


c

| Leiden ID | Majority cell type | Fraction (%) |
|---|---|---|
| 0 | 007 L2/3 IT CTX Glut | 91.2 |
| 1 | 030 L6 CT CTX Glut | 94.6 |
| 2 | 005 L5 IT CTX Glut | 48.0 |
| 3 | 327 Oligo NN | 99.9 |
| 4 | 006 L4/5 IT CTX Glut | 99.0 |
| 5 | 319 Astro-TE NN | 99.1 |
| 6 | 004 L6 IT CTX Glut | 86.9 |
| 7 | 006 L4/5 IT CTX Glut | 26.0 |
| 8 | 334 Microglia NN | 98.6 |
| 9 | 022 L5 ET CTX Glut | 98.7 |
| 10 | 052 Pvalb Gaba | 98.3 |
| 11 | 326 OPC NN | 99.5 |
| 12 | 053 Sst Gaba | 97.9 |
| 13 | 032 L5 NP CTX Glut | 99.9 |
| 14 | 046 Vip Gaba | 80.5 |
| 15 | 049 Lamp5 Gaba | 91.1 |
| 16 | 029 L6b CTX Glut | 85.4 |
| 17 | 327 Oligo NN | 95.0 |
| 18 | Other | 98.4 |
| 19 | Other | 95.4 |
| 20 | 030 L6 CT CTX Glut | 99.9 |
| 21 | Other | 63.3 |
| 22 | 319 Astro-TE NN | 53.5 |
| 23 | 327 Oligo NN | 91.8 |
| 24 | Other | 97.7 |
| 25 | Other | 97.8 |
| 26 | Other | 99.4 |
| 27 | Other | 98.2 |
| 28 | Other | 98.1 |

**Supplementary Fig. 3: Unsupervised Leiden clustering versus MapMyCells**

**(a)** Clusters identified and their IDs via Leiden clustering in the UMAP representation of the snRNA-seq dataset. The nearest neighbors graph was constructed with scanpy's batch corrected K nearest neighbors algorithm scanpy.pp.bbknn(), which was used for Leiden clustering with resolution 1.0.

**(b)** Cell type for each cell was annotated using MapMyCells and labelled in the UMAP representation of the snRNA-seq dataset.

**(c)** For each Leiden cluster, we determined the predominant MapMyCells annotation and calculated the fraction of cells assigned to that cell type. The two approaches showed strong agreement, although Leiden clustering over-segmented certain cell populations. Red text indicates a cell type that was already represented in a larger cluster. Gray shading denotes the corresponding fraction value.

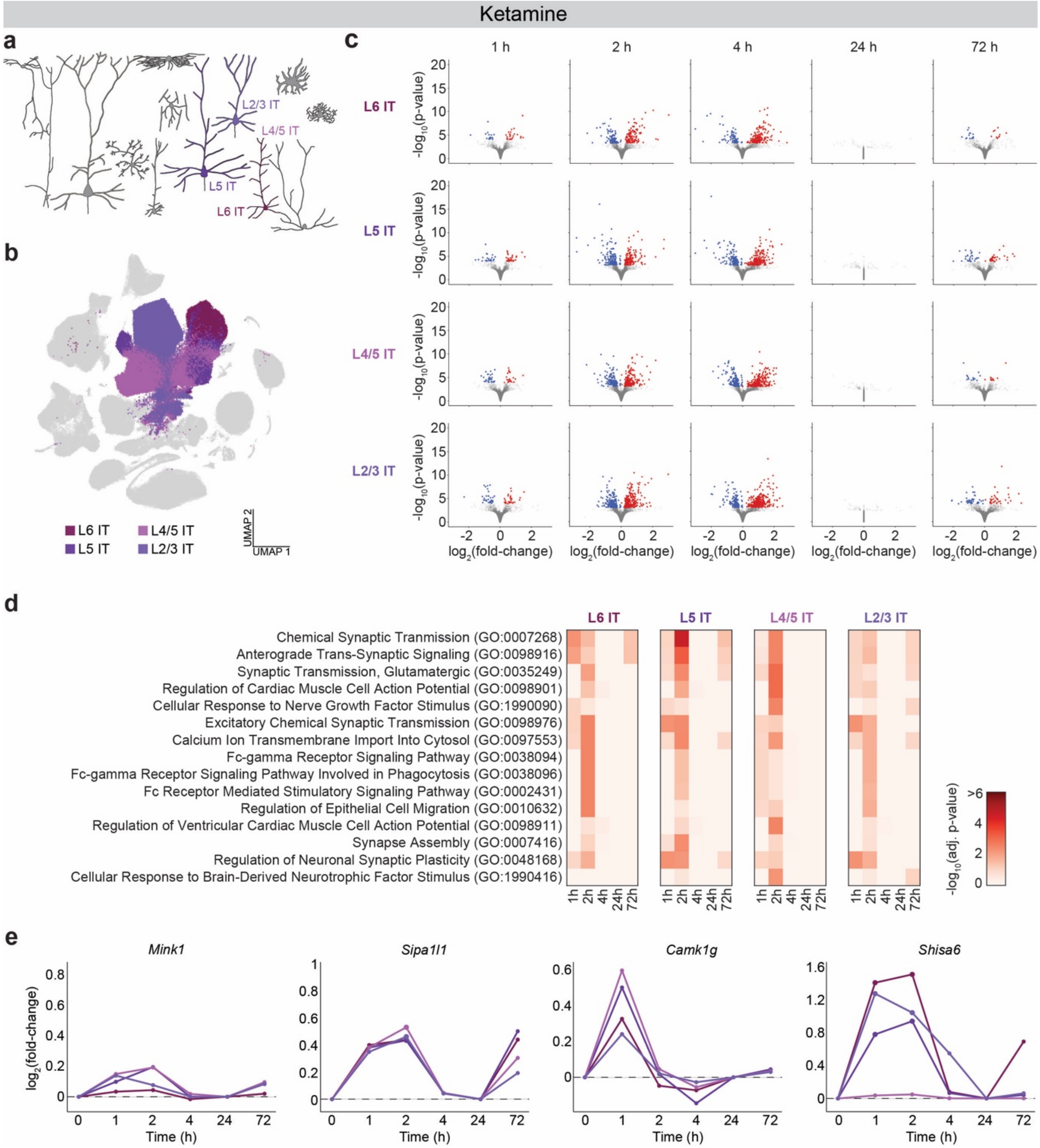


**Supplementary Fig. 4: Ketamine-induced transcriptional changes in frontal cortical IT excitatory neurons**

**(a)** Schematic of the cortical microcircuit highlighting the four IT subtypes of excitatory neurons (L6 IT, L5 IT, L4/5 IT, and L2/3 IT).

**(b)** UMAP representation of the snRNA-seq data set highlighting the four cell types.

**(c)** Differentially expressed genes (DEGs) identified by pseudobulk analysis for each time point for the four cell types. Each circle represents an individual gene. Red, upregulated with FDR < 0.05. Blue, downregulated with FDR < 0.05.

**(d)** Top 15 terms from gene ontology enrichment analysis based on the upregulated DEGs, ranked based on the mean adjusted p-values across time points in the four cell types.

**(e)** Expression levels of specific transcripts in the four cell types following ketamine administration, normalized to controls. The same four genes shown in the main psilocybin figures are presented here for direct comparison between the transcriptional responses evoked by ketamine and psilocybin. Color of the line denotes the cell type.

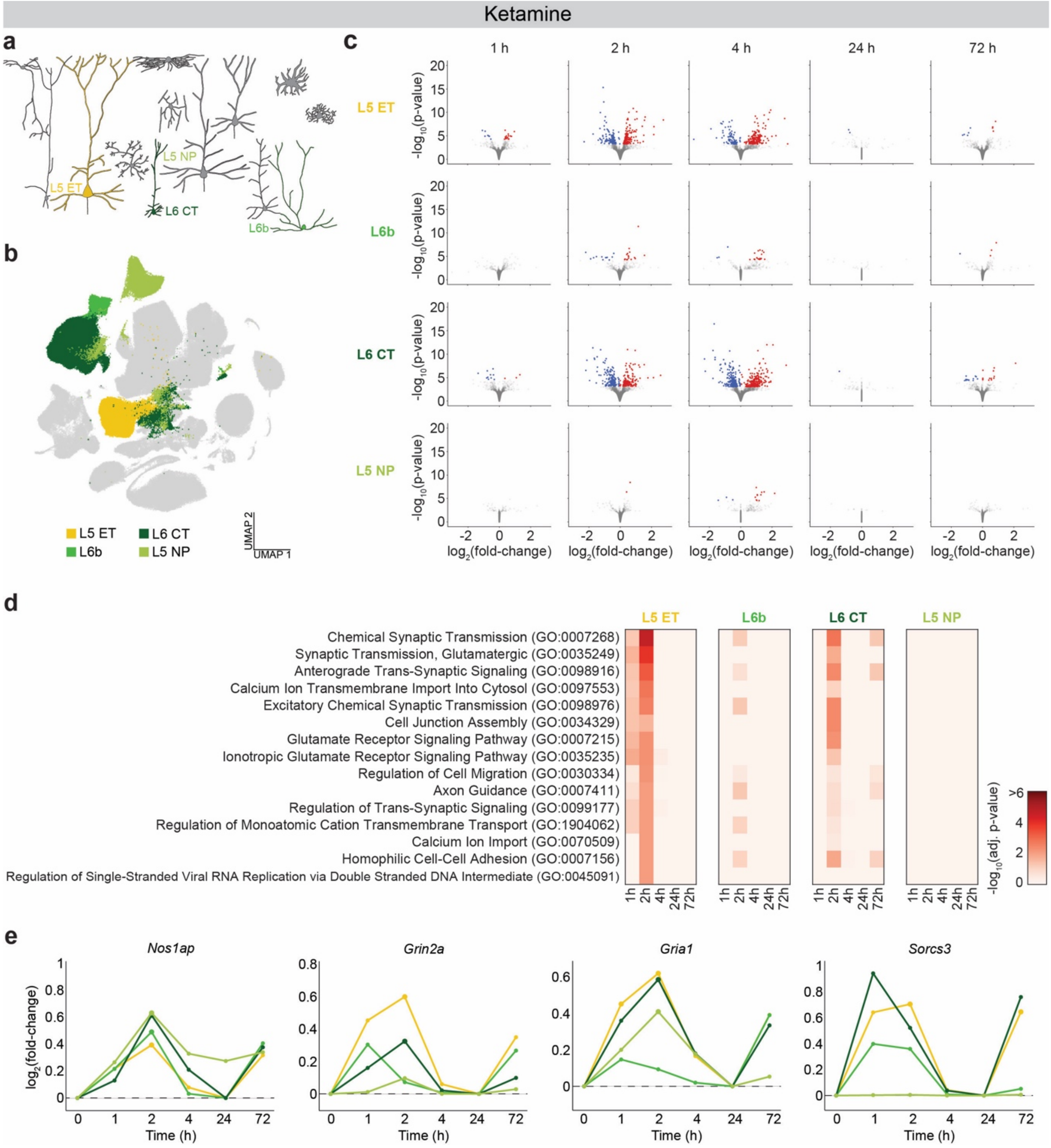


**Supplementary Fig. 5: Ketamine-induced transcriptional changes in other excitatory neuronal subtypes**
**(a)** Schematic of the cortical microcircuit highlighting the four non-IT subtypes of excitatory neurons (L5 ET, L6b, L6 CT, L5 NP).
**(b)** UMAP representation of the snRNA-seq data set highlighting the four cell types.
**(c)** Differentially expressed genes (DEGs) identified by pseudobulk analysis for each time point for the four cell types. Each circle represents an individual gene. Red, upregulated with FDR < 0.05. Blue, downregulated with FDR < 0.05.
**(d)** Top 15 terms from gene ontology enrichment analysis based on the upregulated DEGs, ranked based on the mean adjusted p-values across time points in the four cell types.
**(e)** Expression levels of specific transcripts in the four cell types following ketamine administration, normalized to controls. The same four genes shown in the main psilocybin figures are presented here for direct comparison between the transcriptional responses evoked by ketamine and psilocybin. Color of the line denotes the cell type.

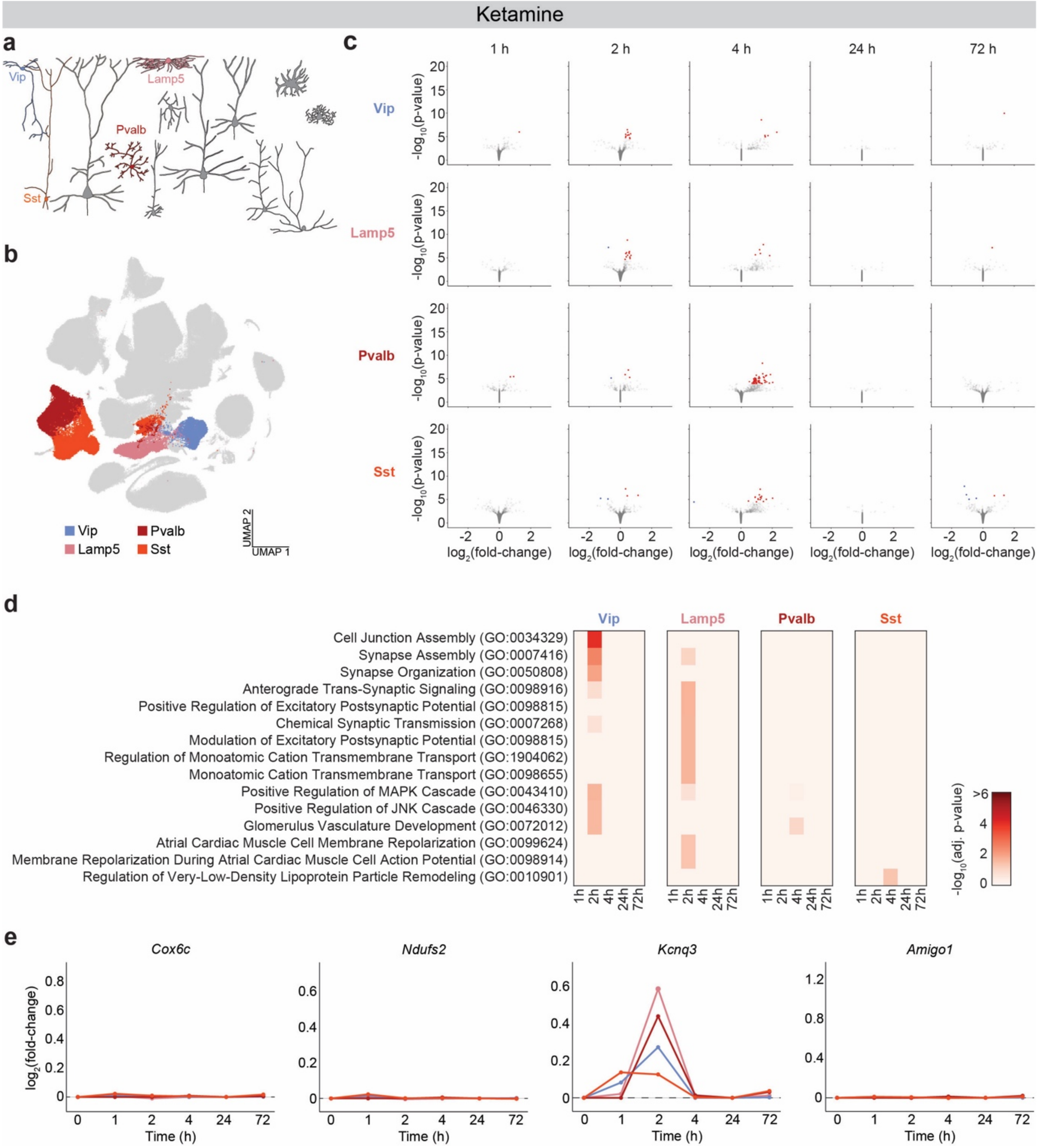


**Supplementary Fig. 6: Ketamine-induced transcriptional changes in frontal cortical GABAergic neurons contribute to mitochondrial function and metabolic processes**

**(a)** Schematic of the cortical microcircuit highlighting the four major classes of GABAergic neurons (Vip, Lamp5, Pvalb, Sst).

**(b)** UMAP representation of the snRNA-seq data set highlighting the four cell types.

**(c)** Differentially expressed genes (DEGs) identified by pseudobulk analysis for each time point for the four cell types. Each circle represents an individual gene. Red, upregulated with FDR < 0.05. Blue, downregulated with FDR < 0.05.

**(d)** Top 15 terms from gene ontology enrichment analysis based on the upregulated DEGs, ranked based on the mean adjusted p-values across time points in the four cell types.

**(e)** Expression levels of specific transcripts in the four cell types following ketamine administration, normalized to controls. The same four genes shown in the main psilocybin figures are presented here for direct comparison between the transcriptional responses evoked by ketamine and psilocybin. Color of the line denotes the cell type.

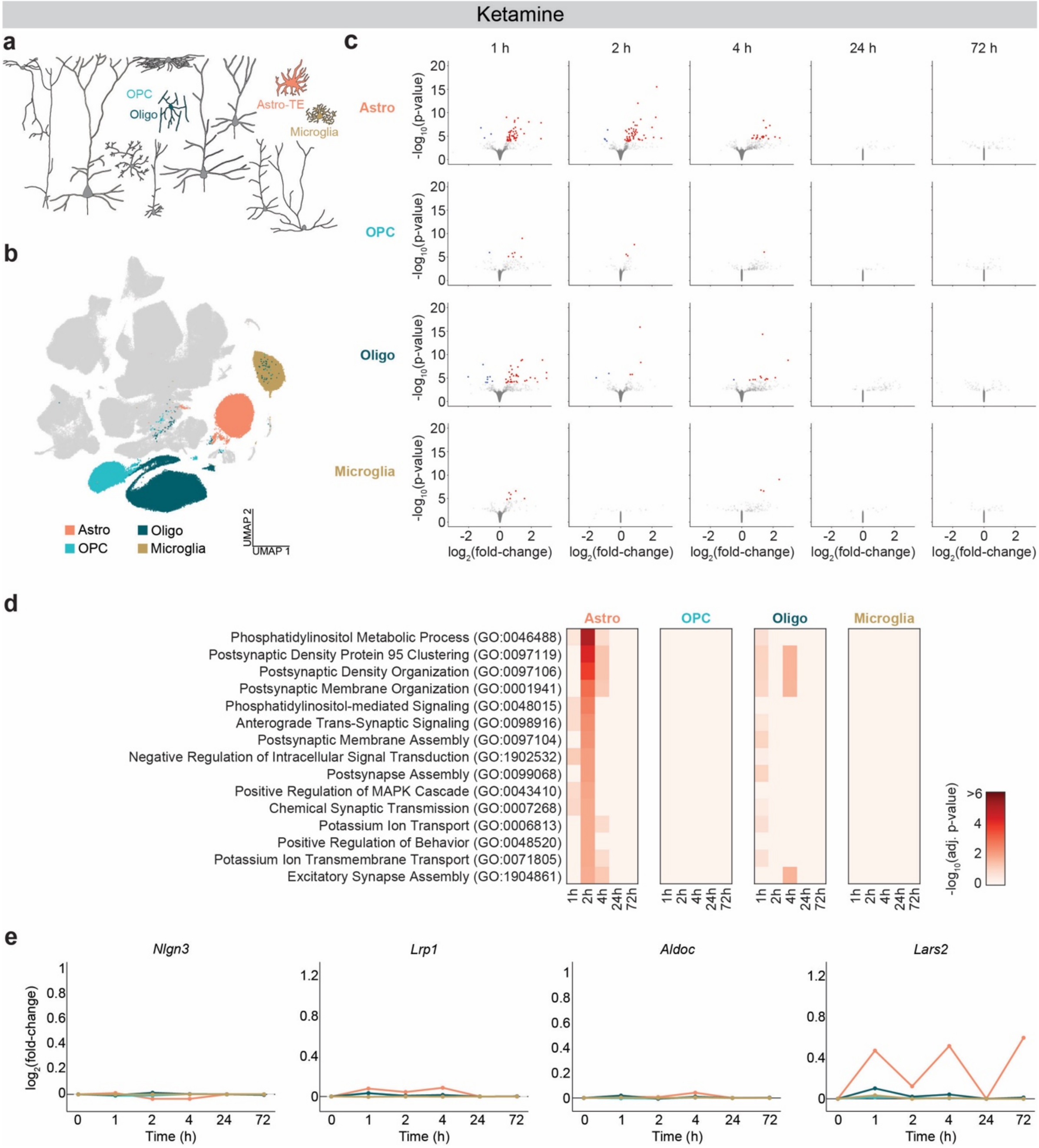


**Supplementary Fig. 7: Ketamine-induced transcriptional changes in non-neuronal cells**

**(a)** Schematic of the cortical microcircuit highlighting the four non-neuronal cell types (Astro, OPC, Oligo, Microglia).

**(b)** UMAP representation of the snRNA-seq data set highlighting the four cell types.

**(c)** Differentially expressed genes (DEGs) identified by pseudobulk analysis for each time point for the four cell types. Each circle represents an individual gene. Red, upregulated with FDR < 0.05. Blue, downregulated with FDR < 0.05.

**(d)** Top 15 terms from gene ontology enrichment analysis based on the upregulated DEGs, ranked based on the mean adjusted p-values across time points in the four cell types.

**(e)** Expression levels of specific transcripts in the four cell types following ketamine administration, normalized to controls. The same four genes shown in the main psilocybin figures are presented here for direct comparison between the transcriptional responses evoked by ketamine and psilocybin. Color of the line denotes the cell type.

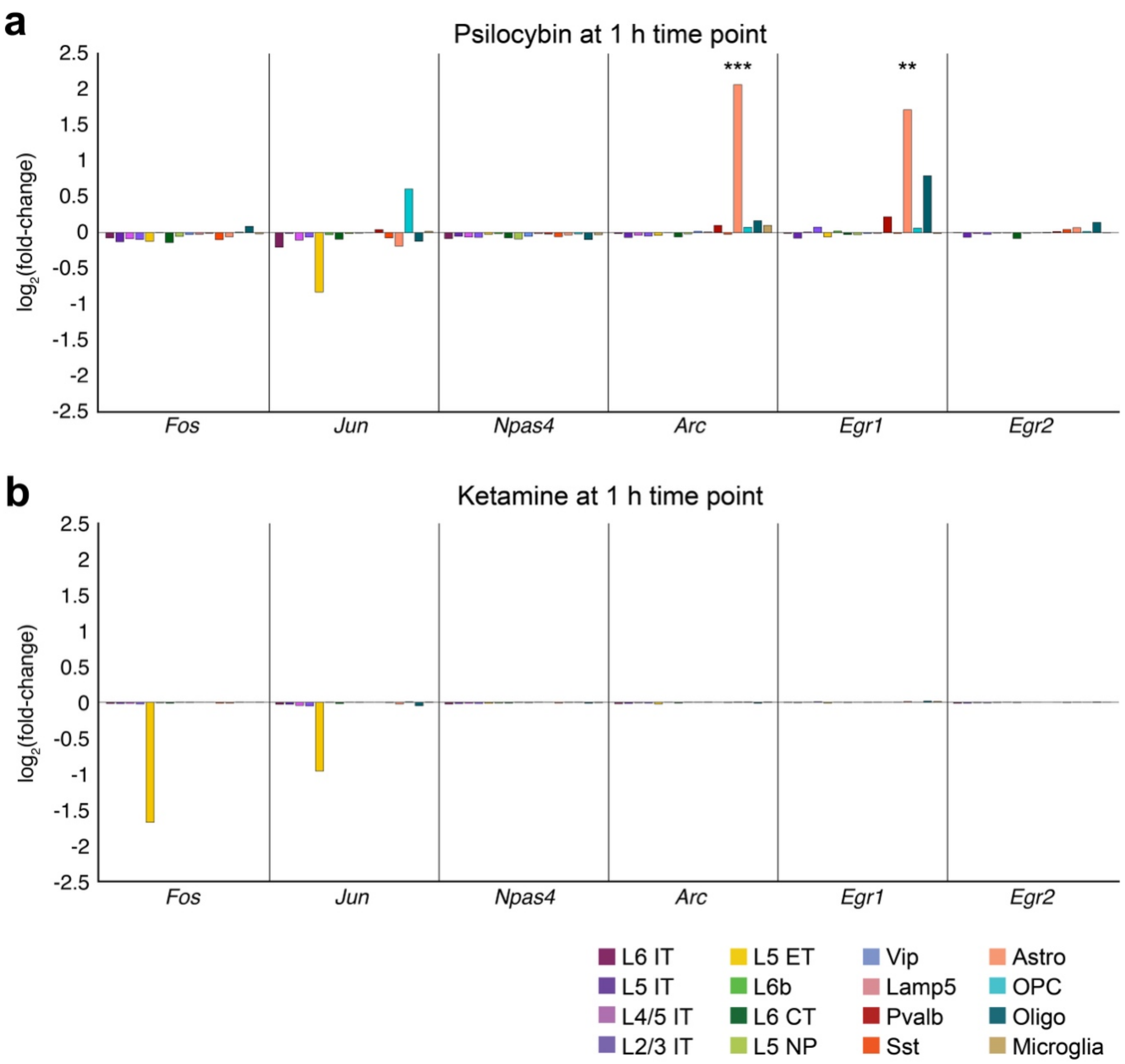


**Supplementary Fig. 8: Immediate early genes**
**(a)** Fold-change in expression for immediate early gene transcripts after psilocybin administration at 1 hour. The fold-change in expression for six Fos, Jun, Npas4, Arc, Egr1, and Egr2 transcripts via pseudobulk analysis for the 16 cell types at 1 hour following psilocybin administration. The color of the bar corresponds to the cell type, using the same color scheme as Fig. 1b. **, adjusted p-value < 0.01. ***, adjusted p-value < 0.001.
**(b)** Similar to (a) for 1 hour following ketamine administration.
